# Using CRISPR/Cas9 to investigate the role of candidate human disease gene orthologs in Ciona

**DOI:** 10.64898/2026.08.10.743552

**Authors:** Sabrina A. Hernandez, Christopher J. Johnson, Alberto Stolfi

## Abstract

The tunicate *Ciona robusta* offers a tractable non-vertebrate chordate model for probing gene function via tissue-specific, CRISPR/Cas9-mediated mutagenesis in F0. Building on Arcadia Science’s Zoogle platform, which identifies and ranks orthologs of human genes from various non-traditional model organisms, we carried out a pilot project to probe the developmental roles of three notochord- and endoderm-expressed candidate orthologs of human disease genes (*Fcho*, *Pgm3*, and *Nckap1*) alongside a fourth gene (*Plastin*) implicated in papilla cell elongation. This preprint compiles and updates a series of research project milestones previously posted episodically on Zenodo. Here we summarize the full results and our conclusion about this pilot project. Using CRISPR/Cas9, we found that tissue-specific knockout of *Pgm3* and, to a lesser extent, *Fcho* caused significant defects in larval tail elongation. Separately, CRISPR knockout of *Plastin*, an actin-bundling gene expressed throughout the sensory-adhesive papillae of the larva, caused a subtle reduction in papilla cell elongation when combined as a duoble knockout with another actin-bundling protein-encoding gene, *Villin*. These results identify *Pgm3* as the most promising candidate for further development as a Ciona-based model of human disease and demonstrate the utility of tissue-specific CRISPR screening for prioritizing candidate disease gene orthologs identified through comparative genomics platforms like Zoogle.

## INTRODUCTION

The tunicate *Ciona robusta* (formally *intestinalis* Type A) serves as a convenient chordate model for investigating gene function during embryogenesis (Satoh, 2013). Its simplified body plan, rapid development, and experimental tractability make it an attractive system for exploring fundamental or potentially chordate-specific mechanisms of gene regulation and cellular morphogenesis (Bernadskaya and Christiaen, 2016). CRISPR/Cas9-mediated mutagenesis in *Ciona* has enabled targeted disruption of single genes with high efficiency in F₀ embryos, supporting both basic research and pedagogical applications (Johnson et al., 2023; Pennati et al., 2024).

With support from Arcadia Science, our group has used CRISPR/Cas9 in *Ciona* to investigate the developmental roles of candidate genes identified by Arcadia Science’s Zoogle platform (https://zoogle.arcadiascience.com/about) as *Ciona* orthologs of human disease-associated genes (Chou et al., 2025; Chou and Sun, 2025b). This report combines four installments of that ongoing project, previously published on Zenodo (Hernandez et al., 2025; Hernandez and Stolfi, 2026; Johnson et al., 2025; Johnson and Stolfi, 2025). The first three describe our investigation of three notochord- and endoderm-expressed genes (*Fcho*, *Pgm3*, and *Nckap),* from sgRNA design through phenotypic analysis of tail morphogenesis. The fourth describes a related, parallel investigation, inspired by an Arcadia Science collaboration with David Booth’s group at UCSF in the choanoflagellate *Salpingoeca rosetta* (Chou and Sun, 2025a), of the gene *Plastin* and its role in elongation of the sensory-adhesive papilla cells of the *Ciona* larva.

Targeting this small set of genes was intended as a pilot project to identify potential candidate genes for which *Ciona* might serve as a useful human disease model. The first step was to identify loss-of-function phenotypes by tissue-specific CRISPR. Here we describe in detail our CRISPR approach and report the initial characterization of these knockouts, which might inform future plans for using *Ciona* as a model to study the associated human disease genes. Of note, both *Fcho* and *Pgm3* appear to be required for proper morphogenesis of the *Ciona* larval tail, while *Plastin* might be required in combination with another gene, *Villin,* for proper cell elongation.

## METHODS, RESULTS, AND DISCUSSION

### Fcho, Pgm3, and Nckap1 identified as target candidate genes

*Fcho*, *Pgm3*, and *Nckap1* were identified as *Ciona* orthologs of human disease-associated genes and intriguing candidates for further investigation by Arcadia Science’s Zoogle platform (**Figure 1A**)(Chou et al., 2025; Chou and Sun, 2025b). In vertebrates, Fcho1 (FCH And Mu Domain Containing Endocytic Adaptor 1) encodes an endocytic adaptor involved in clathrin-mediated vesicle formation (Calzoni et al., 2019; Henne et al., 2010; Łyszkiewicz et al., 2020). Pgm3 (Phosphoacetylglucosamine mutase 3) participates in carbohydrate metabolism and glycosylation pathways critical for extracellular matrix integrity (Greig et al., 2007; Stray-Pedersen et al., 2014; Winslow et al., 2022; Yang et al., 2024). Nckap1l (Nck-Associated Protein 1-Like) is part of the WAVE regulatory complex, linking signaling pathways to actin cytoskeleton remodeling (Castro et al., 2020; Cook et al., 2020; Park et al., 2008). Disruption of these genes in humans is associated with diverse developmental and immune disorders, but we lack a more detailed tissue-level understanding of their cellular functions.

**Figure 1.**
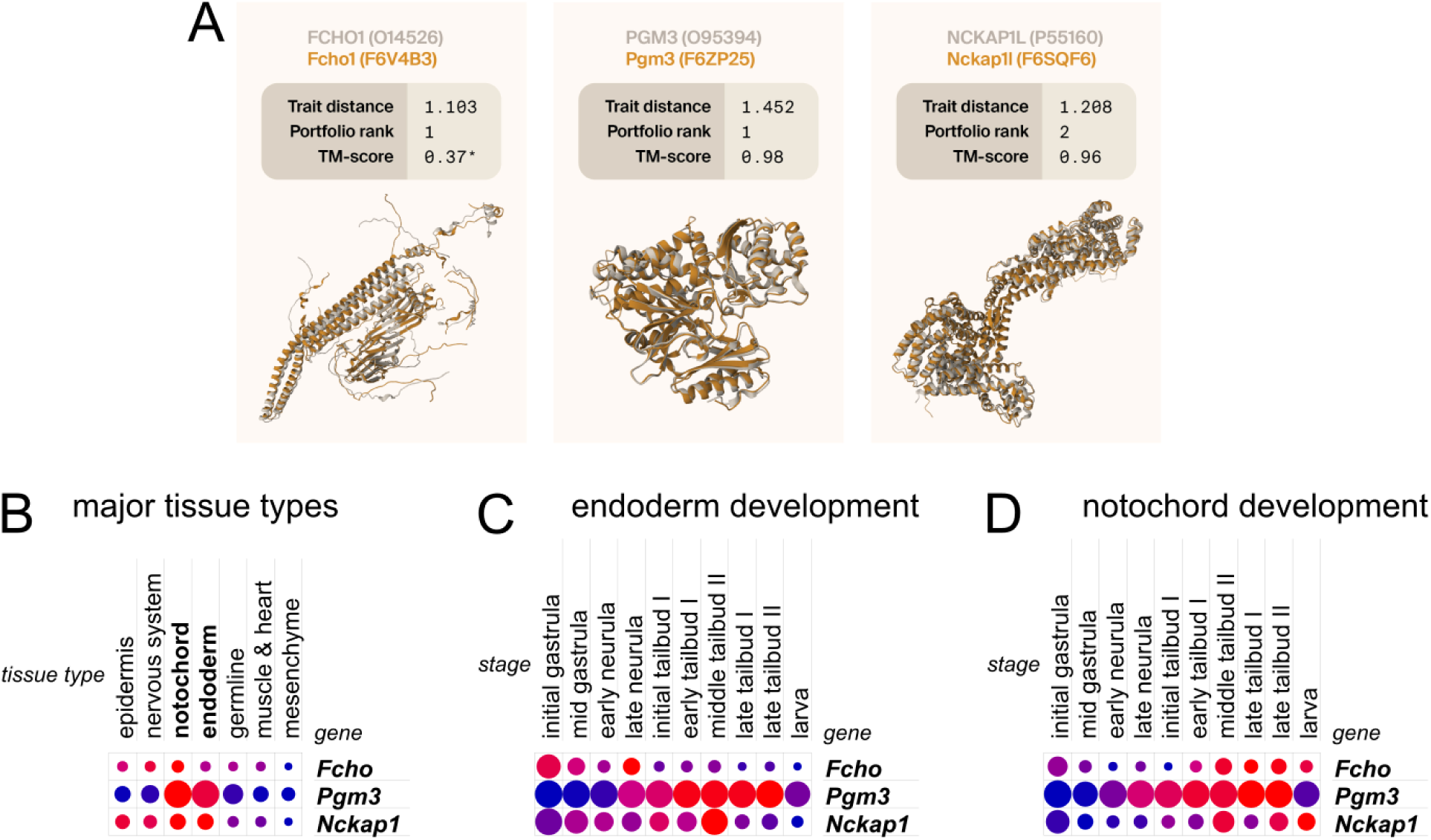
A) Comparison of human disease proteins FCHO1, PGM3, and NCKAP1L and their *Ciona* orthologs, figure adapted from Chou and Sun 2025b, courtesy of Arcadia (Chou and Sun, 2025b). Trait distance and portfolio ranks are generated by Zoogle, with detailed methods explained in Chou et al. 2025. “Portfolio rank” is the relative rank of the *Ciona* ortholog compared to all other species’ orthologs (and sometimes paralogs). TM-score is a standard measure of protein structural similarity. See Chou and Sun 2025b for more detailed information about protein structure predictions. B) Single-cell RNAseq (scRNAseq) expression plot comparing the three genes’ expression across different tissue clusters in the whole developmental dataset from Cao et al. 2019, as visualized by Single Cell Portal. C) Similar plot comparing expression in the endoderm across developmental stages. D) Similar plot as in C, but for notochord expression across developmental stages. For plots B-D, the relative size of the dot is proportional to % cells expressing the gene, while the blue-red gradient indicates scaled mean expression from 0 (blue) to 1 (red).

As highlighted previously by both bulk and single-cell RNAseq (scRNAseq), all three appear to be expressed in the notochord and endoderm during *Ciona* embryonic development (**Figure 1B-D**)(Cao et al., 2019; Chou and Sun, 2025b). Only *Pgm3* expression has been characterized by *in situ* hybridization, showing strong and specific upregulation in the endoderm and notochord (Reeves et al., 2017). Hence, we started focusing on a plan to knock these genes out in endoderm and notochord cell lineages and assay the effects on the proper morphogenesis of these cells and surrounding tissues during embryogenesis.

### Designing sgRNAs to target *Fcho, Pgm3,* and *Nckap1*

For each target gene, three single-chain guide RNAs (sgRNAs) were selected (**Table 1**) using the CRISPOR website (https://crispor.gi.ucsc.edu/) (Concordet and Haeussler, 2018), following our established *Ciona*-specific strategies that have been previously published (Popsuj et al., 2024). CRISPOR is ideal for researchers working on *Ciona* due to its incorporation of single nucleotide polymorphism (SNP) data, allowing us to avoid potentially polymorphic target sites; *Ciona* populations are known to harbor extreme genetic diversity (Nydam and Harrison, 2010), and any mismatches between the sgRNA and the target sequence in the actual specimens used in the lab might result in low mutagenesis efficacies.

**Table 1.** List of novel sgRNAs designed to target *Fcho, Pgm3,* and *Nckap1*, and their efficacy and specificity scores as predicted by CRISPOR. Asterisks denote “false” cases of low specificity due to confirmed genome assembly errors.

| sgRNA | Doench '16 | Doench-RuleSet3 | MIT specificity |
| --- | --- | --- | --- |
| Fcho 1.34 | 57 | 23 | 100 |
| Fcho 4.63 | 59 | 33 | 100 |
| Fcho 5.112 | 63 | -25 | 99 |
| Nckap1 2.74 | 68 | 63 | 100 |
| Nckap1 7.36 | 66 | 43 | 100 |
| Nckap1 8.103 | 60 | 71 | 100 |
| Pgm3 1.83 | 72 | 81 | 33* |
| Pgm3 2.27 | 60 | 24 | 50* |
| Pgm3 3.31 | 59 | 61 | 100 |

To search for potential sgRNAs, we entered each gene’s exons one at a time, starting from exon 1 upwards, until we obtained sgRNAs with high enough predicted efficacies and specificities. High efficacy is typically predicted by high “Doench ’16” score (Doench et al., 2016), specificity by an “MIT specificity” score >99 (Hsu et al., 2013). We have previously shown correlation between high, actual efficacy and predicted Doench ’16 efficacy scores over 50. Thus, our informal predicted efficacy (Doench ’16) score cutoff is typically ∼50. A new predictive algorithm has been recently incorporated into CRISPOR, named “Doench RuleSet 3” (RS3)(DeWeirdt et al., 2022). There is some evidence that RS3 might be modestly better at predicting effective sgRNAs for CRISPR in *Ciona* (Popsuj et al., 2026), though this remains to be rigorously shown.

To ensure specificity, we aimed for MIT specificity scores >99. As the *Ciona robusta* genome is quite small and compact (∼120 Mbp), scores of 99 and 100 are quite common and would rule out most potential off-target effects. We have previously detected a low (∼5%) mutagenesis rate at a site with perfect sgRNA seed-target match but a non-canonical/alternative protospacer-adjacent motif (PAM)(Popsuj et al., 2024). However, CRISPOR did not allow us to automatically screen for those.

Candidate sgRNAs were chosen to maximize predicted mutagenesis efficiency while minimizing potential off-target activity. We also prioritized targeting sequences encoding the N-terminal region of the resulting protein, to maximize the likelihood of generating null alleles. We must balance the likelihood that a null, or “near-null” allele will result from our CRISPR-mediated indels, with the chance that cutting too close to the start codon will allow for translation of a mostly functional protein from a new, alternate start codon further into the coding sequence.

For Pgm3, we noticed abnormally low MIT specificity scores for sgRNAs targeting the first two exons, but we ascertained that this was due to older genome assembly errors that had erroneously duplicated those Pgm3 sequences. We were able to confirm that these earlier assembly errors were not present in the latest updates “HT” version of the Ciona robusta genome assembly (Satou et al., 2019), thus restoring our confidence in the specificity of these sgRNAs. Indeed, all the predicted off-targets flagged initially were in the incorrect assembly duplicate sequences.

### Validating sgRNAs

All sgRNAs were tested according our established protocol (<u>link to protocols.io</u>)(**Figure 2**)(Popsuj et al., 2024). Briefly, each sgRNA is expressed using the U6 small RNA promoter, from plasmid co-electroporated into synchronized *Ciona* zygotes along with *Eef1a>Cas9* plasmid. In this setup, Cas9 and the sgRNA are expressed in most cells. Larvae are then collected after hatching, comprising pools of hundreds or even thousands of F_0_ CRISPants. Genomic DNA is extracted from each pool (representing a single sgRNA test), and amplicons centered on the sgRNA target site are obtained by PCR from the resulting genomic DNA. Amplicons (150-450 bp in size) are then sequenced by Illumina sequencing by Genewiz/Azenta, which provides automated indel analysis including estimated mutagenesis efficacy rate. Often the efficacy percentages presented are only approximated due to confounding naturally occurring SNPs and indels. This is because of the high genetic diversity in the wild *Ciona* populations we use for our studies.

**Figure 2.**
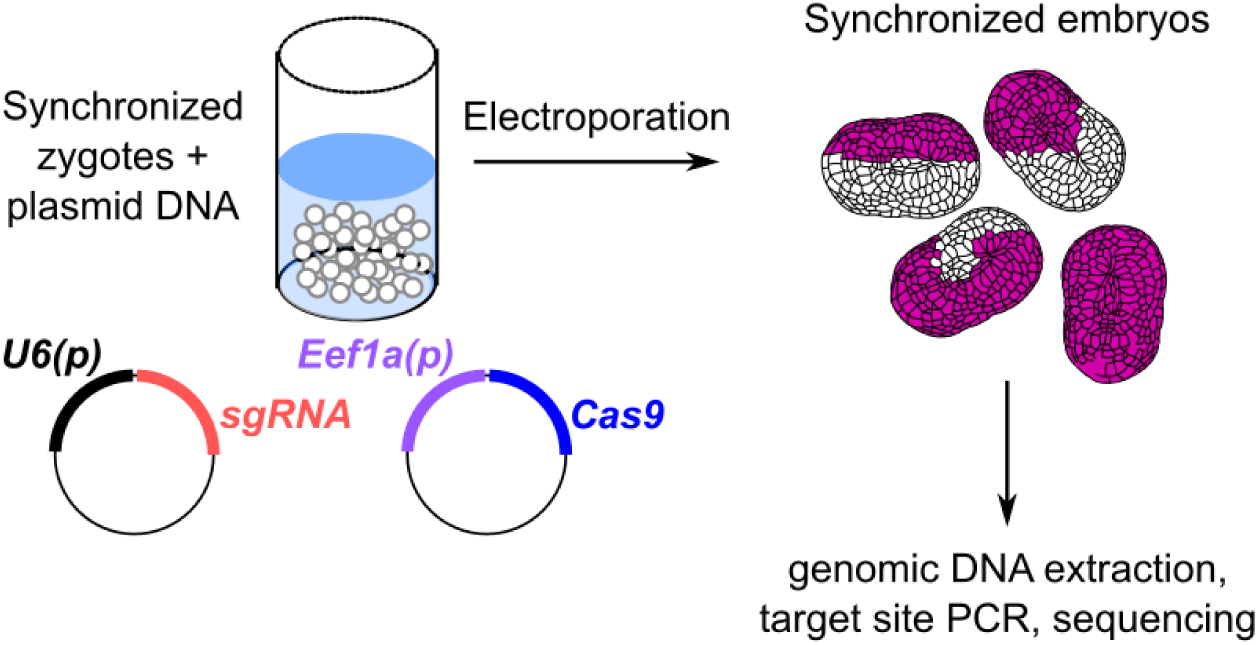
Diagram depicting our sgRNA validation workflow. Synchronized Ciona robusta zygotes (1-cell-stage embryos) are electroporated with plasmid DNAs to drive near-ubiquitous expression of an sgRNA and Cas9. The sgRNA is transcribed by RNA polymerase III via the small RNA promoter, while Cas9 expression is driven by the Eef1a promoter. Co-expression is denoted by purple coloring of cells. Mosaic uptake and/or retention of the plasmids results in some cells that do not express either. Embryos are then left to grow (typically to the larval stage) before pooling and genomic DNA extraction and downstream PCR and sequencing of target site amplicons (see text and linked protocol for details).

For *Fcho,* we tested three sgRNAs targeting exons 1, 3, or 5, surrounding the region that encodes a conserved FCH domain near the N terminus (**Figure 3**). The most efficacious sgRNAs by indel analysis were sgRNAs 1.34 and 5.112. The efficacy of sgRNA 1.34 was measured at 43%, which is on the higher end of the range of efficacies that we have measured for other sgRNAs in our lab. For sgRNA 5.112, an exact efficacy percentage was not obtained due to a confounding natural SNP, but it was well over 15%. Comparing to 1.34, it was likely to be closer to 30-40%. As such, these two sgRNAs were selected for future use, and sgRNA 4.63 was all but abandoned.

**Figure 3.**
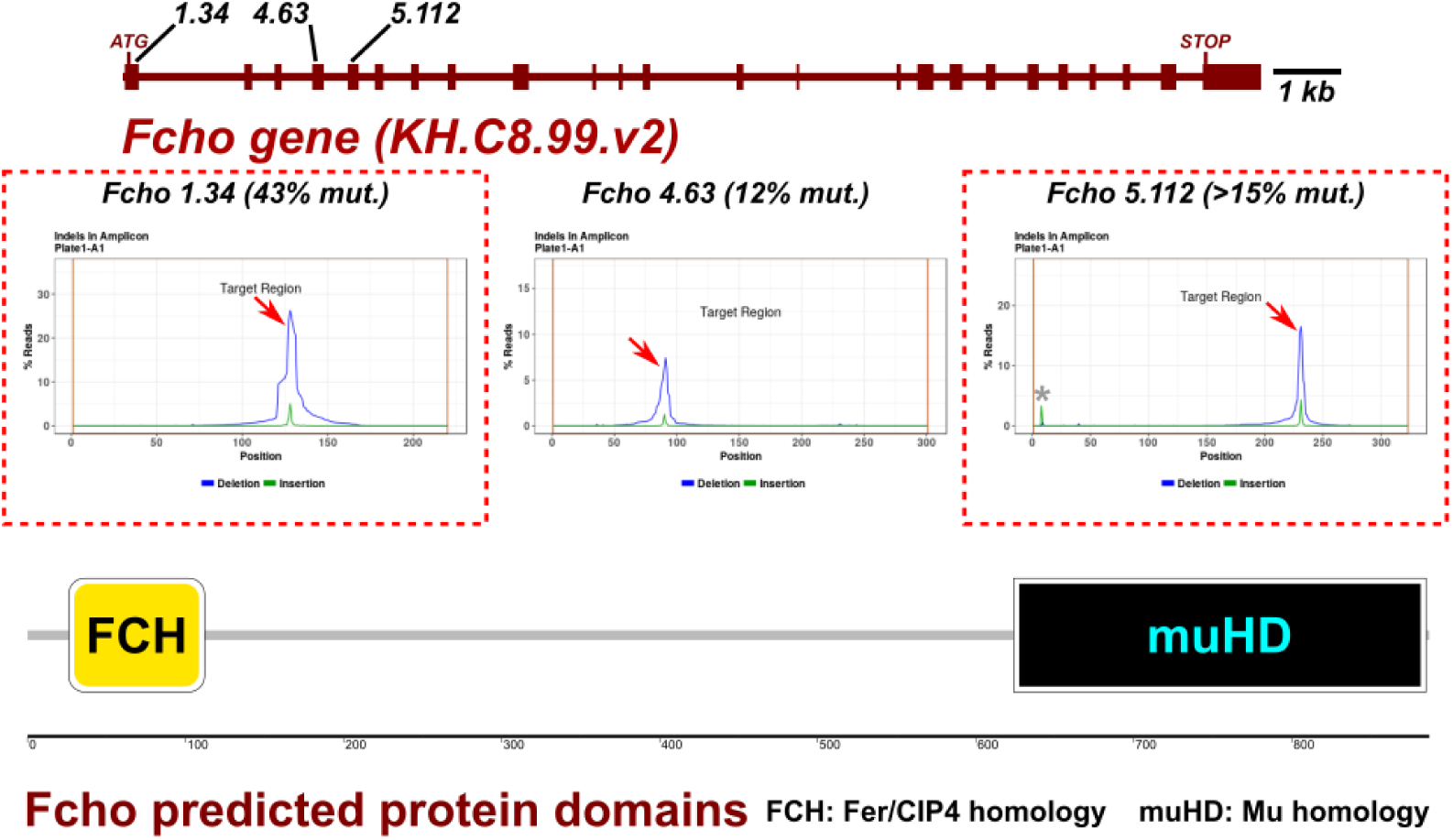
*Fcho* sgRNA validation results

When used in combination, sgRNAs 1.34 and 5.112 are also predicted to delete the entire FCH domain at the N terminus, which would further increase the odds of producing a null allele.

For *Pgm3,* sgRNAs 1.83 and 2.27 had the highest efficacies (**Figure 4**). Again, the exact efficacy for 1.83 was not measured due to a naturally occurring indel in the amplicon region, but was well over 12%. By looking at the height of the major CRISPR-induced deletion peak, its efficacy appeared roughly equivalent to that of sgRNA 2.27, which was measured at 23%. Both of these sgRNAs target the first two overlapping phosphoglucomutase/phosphomannomutase (PGM/PMM) domains, suggesting that using them in combination will likely produce null alleles. We therefore went ahead and selected these two for further use.

**Figure 4.**
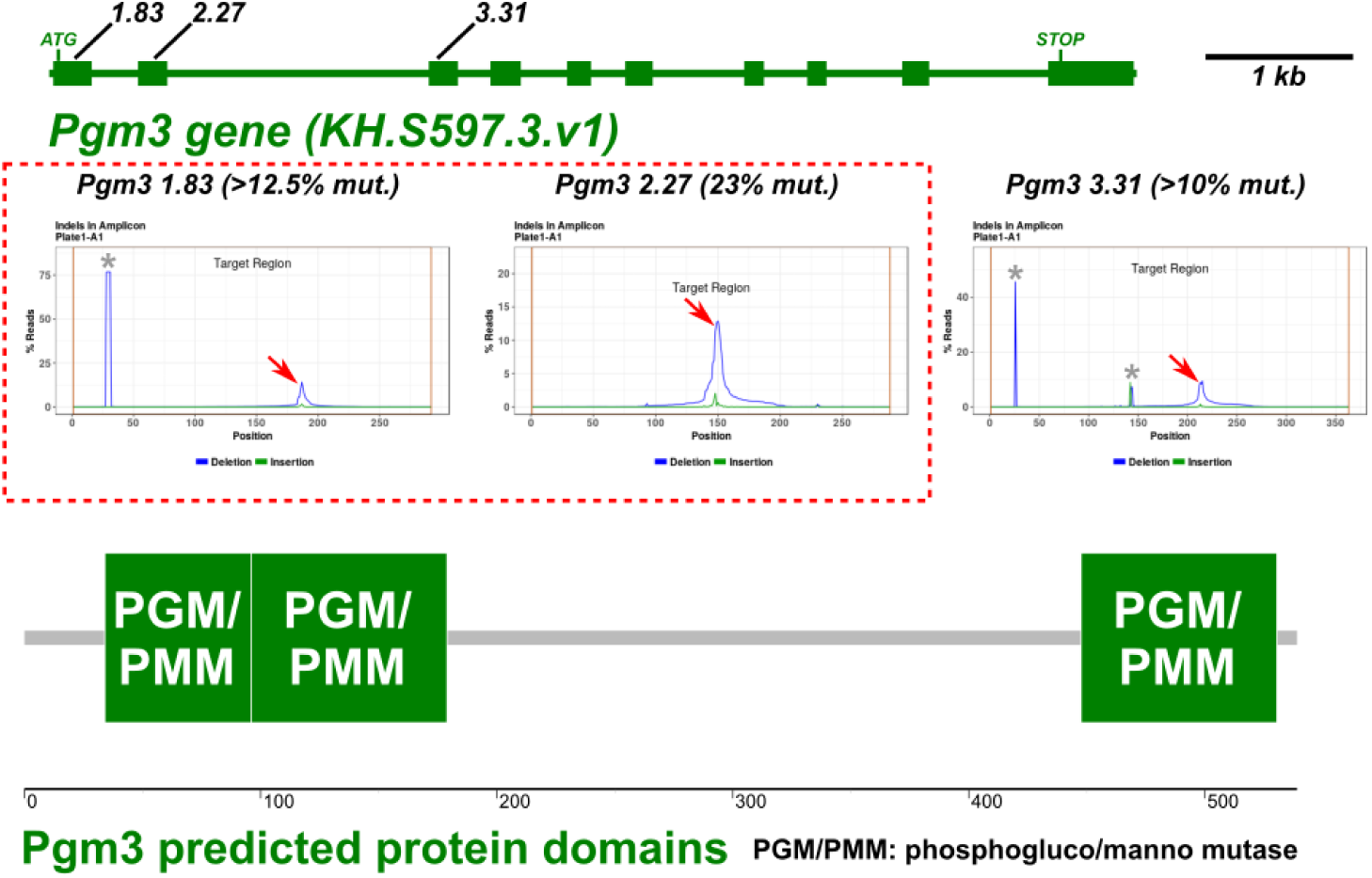
*Pgm3* sgRNA validation results

Finally, for *Nckap1* (**Figure 5**), the presence of extensive natural indels and SNPs prevented an exact measurement of the efficacy of any of our three sgRNAs, but based on comparisons to the *Pgm3-*targeting sgRNAs, we believe that 2.74 and 8.103 were also in the 20-30% range. These target the 2^nd^ and 8^th^ exons of *Nckap1,* respectively. Unlike Fcho or Pgm3, the Nckap1 protein does not have any conserved domains other than the entire protein representing its own domain family. As such, it is not clear what parts of this large protein might be key for its functions, but using the above sgRNAs in combination, a large deletion at the N terminus should likely result in null alleles.

**Figure 5.**
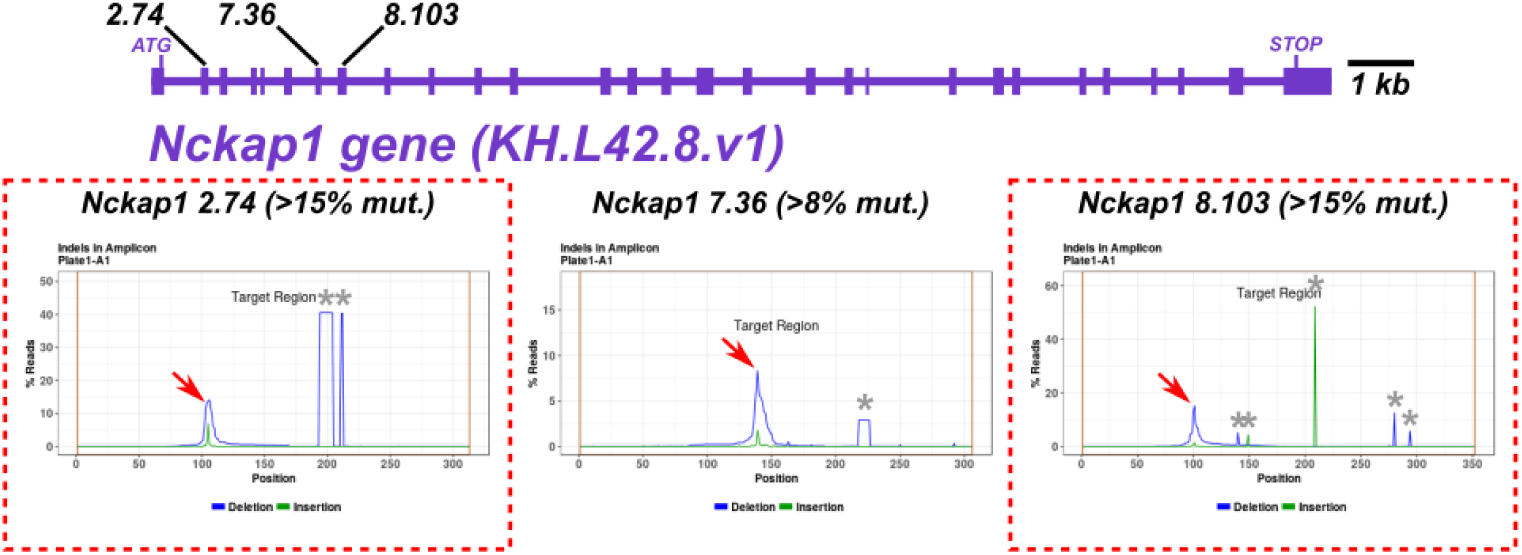
*Nckap1* sgRNA validation results

### Phenotyping *Fcho, Pgm3,* and *Nckap1* CRISPR knockouts in endoderm and notochord

Because all three targeted genes are expressed in the *Ciona* notochord during embryogenesis (**Figure 1B.D**) we first asked if any are required for *Ciona* notochord convergent extension, an essential process that occurs early during notochord development (**Figure 6**). This process involves the mediolateral polarization and migration of 40 post-mitotic cells that form two rows of 20 cells on either side of the midline in the developing embryo (Di Gregorio, 2020). These two rows of cells then intercalate to form a single row of cells. Eventually, through additional steps of elongation, vacuolization, and tubulogenesis, the notochord transforms into an extracellular fluid-filled tube that serves as a hydrostatic skeleton for the swimming *Ciona* larvae (Jiang and Smith, 2007).

**Figure 6:**
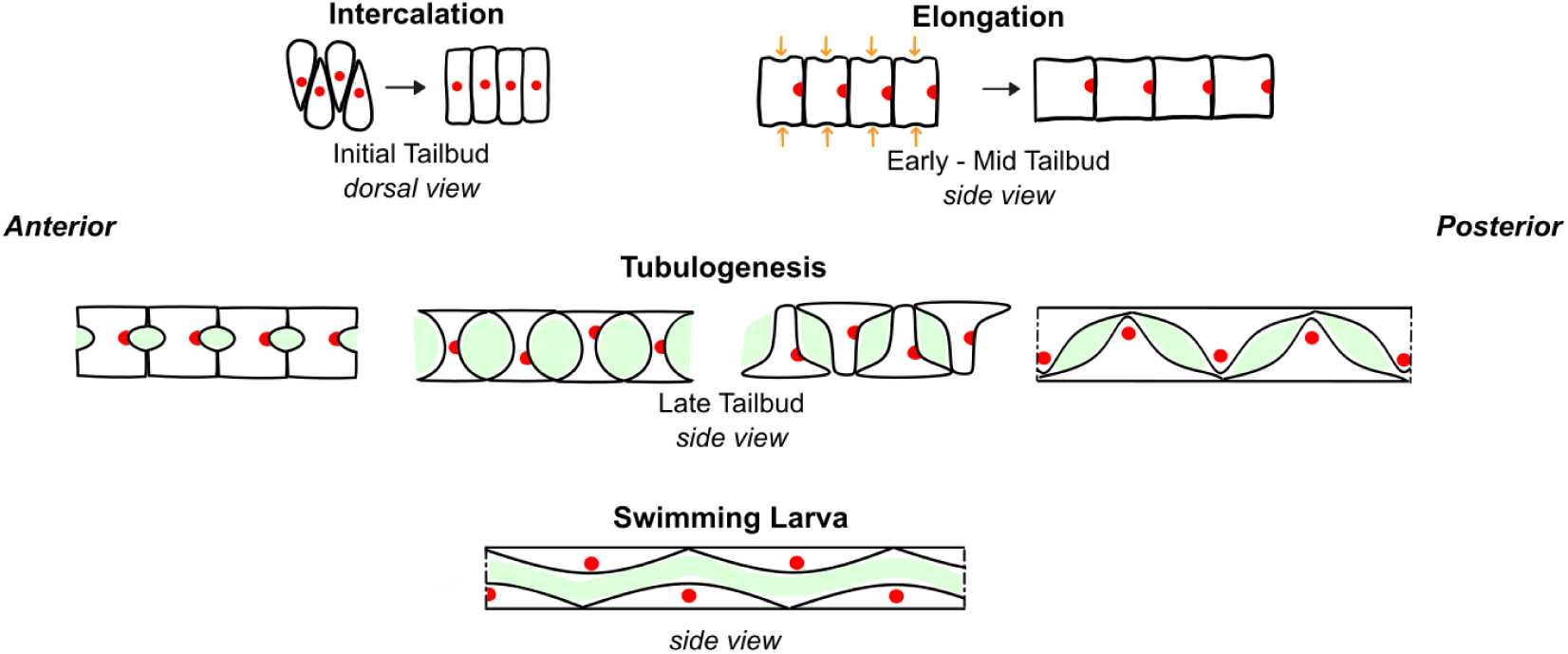
schematic of notochord morphogenesis in Ciona during embryogenesis.

With validated sgRNAs in hand, we used specific combinations of the most effective sgRNAs for each gene to target each one as follows. To knock out *Pgm3,* we used the combination of sgRNAs 1.83 and 2.27. For *Fcho,* we used the 1.34 and 5.112 sgRNA combination, and for *Nckap1*, we combined sgRNAs 2.74 and 8.103. These were previously identified as having the highest efficacies in creating insertion/deletion in our desired target regions (Johnson et al.). These data are summarized in **Table 2** below.

**Table 2:** Estimated efficacies of selected sgRNAs. See validation above for details.

| sgRNA | Mutation efficacy (indel rate) |
| --- | --- |
| Pgm3.1.83 | 23% |
| Pgm3.2.27 | >12.5% |
| Fcho.1.34 | 43% |
| Fcho.5.112 | >15% |
| Nckap1.2.74 | >15% |
| Nckap1.8.103 | >15% |

To assess the potential role of these genes in notochord cell intercalation, we performed CRISPR/Cas9-mediated mutagenesis in the *Ciona* early cell lineages that give rise to the notochord and endoderm. We used the dechorionation and electroporation protocol previously described by Christiaen et al (Christiaen et al., 2009a, b). Synchronized zygotes were electroporated with gene-specific combinations of plasmids encoding the sgRNAs described in **Table 2**, together with the *Foxa.a>Cas9::Geminin^N-terminus^* plasmid to induce mutations early in endoderm and notochord lineages as previously described (Di Gregorio et al., 2001; Popsuj et al., 2023). We allowed the embryos to develop to the mid tailbud I stage (Hotta Stage 21, roughly 9.5 hours post-fertilization when reared at 20°C) when intercalation is complete in normally developed embryos. The fluorescent reporter plasmids *Brachyury>CD4::GFP* (encoding cell membrane-bound GFP), *Brachyury>Unc-76::GFP* (encoding GFP tagged with a portion of the Unc-76 protein from *C. elegans,* which promotes its cytoplasmic localization), and/or *Brachyury>H2B::mCherry* (encoding histone-fused mCherry) were also used to visualize the notochord cell shape and nuclei, respectively (Corbo et al., 1997). A control experiment was performed using the same plasmids, using a *U6>control* sgRNA plasmid targeting no *Ciona* sequence instead of the sgRNA plasmid combination for each gene. Per 700 µl of electroporation volume, we used 80 µg of sgRNA plasmid (40 µg of each when combined), 35 µg of Cas9 plasmid, and 40 µg of each green or red reporter plasmid.

Embryos in control and knockout conditions were imaged using an inverted epifluorescence microscope and scored as either normal intercalation, minor defect in intercalation (1-2 cells out of midline) and severe defect in intercalation (3+ cells out of midline). Examples of normal and severe defects in three different conditions are shown in **Figure 7**. Experiments were performed in three separate biological replicates using different batches of dissected gametes, with 75 individuals scored per condition and replicate. Raw images and scoring data for *Pgm3* and *Fcho* CRISPR embryos (but not *Nckap1* CRISPR) can be found here: https://zenodo.org/records/17857562

**Figure 7:**
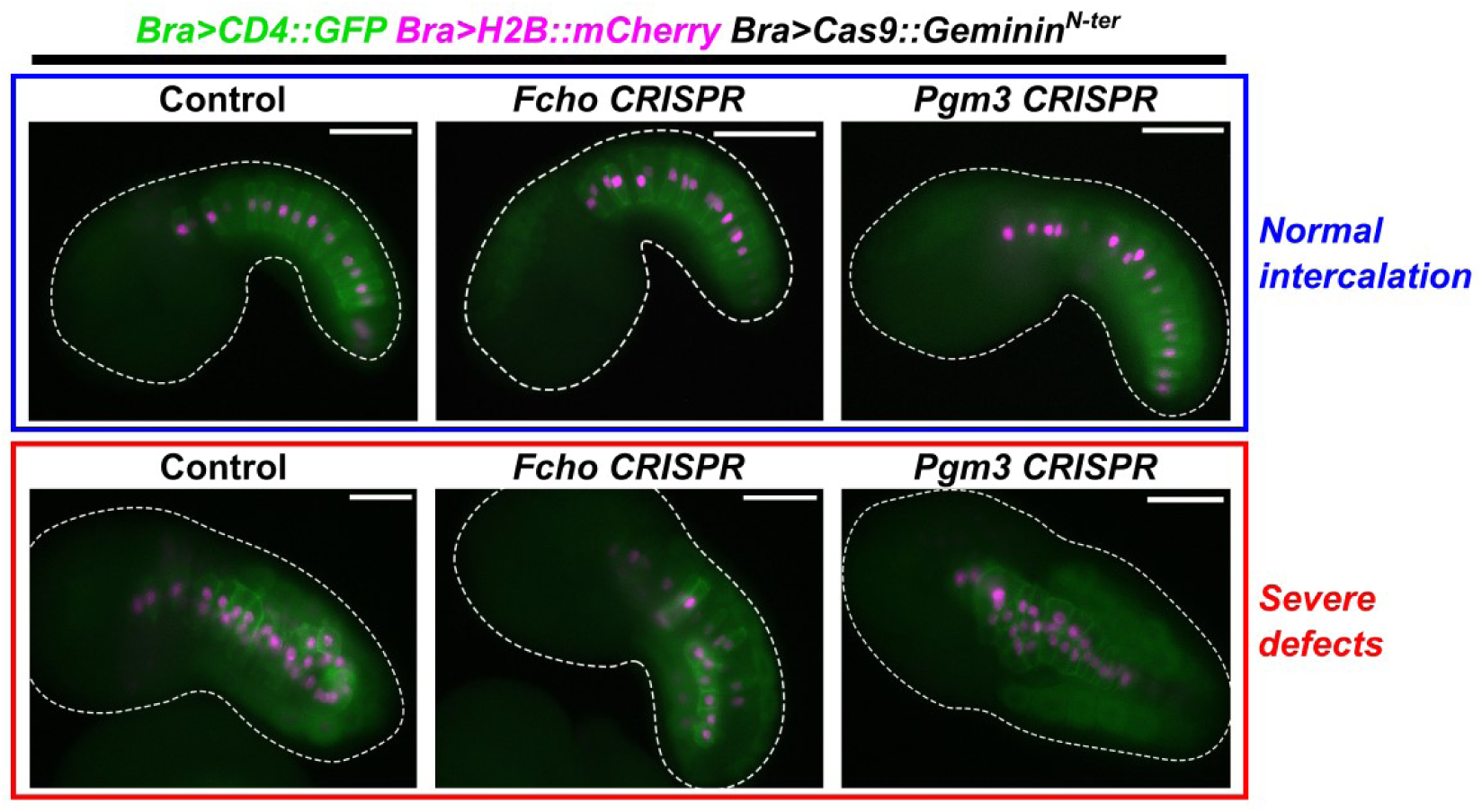
Representative epifluorescence microscopy images of early tailbud embryos showing examples scored as “normal” intercalation (top) or those with severe intercalation defects (bottom). Control = negative control CRISPR condition. Dashed outlines indicate embryo outline. Anterior always to the left or top/bottom-left. Dorsal always over ventral. All scale bars = 50 µm. See text for details about type and amount of plasmids electroporated.

Three replicates of the *Pgm3* knockout were performed, and the scoring data are summarized in **Figure 8**. In the first replicate, we observed severe notochord intercalation defects in 51% of embryos in the *Pgm3* knockout condition, compared to 23% of negative control embryos. This initially suggested that Pgm3 could play a role in proper notochord intercalation. We performed an additional two replicates of this experiment. However, in these additional experiments, we observed roughly equal percentage of defects in both control and knockout conditions (∼50% and ∼30% defects in replicates two and three respectively). Since we could not replicate our initial results showing an increase in intercalation defects in *Pgm3* knockout condition compared to the control, we cannot conclude that *Pgm3* is vital for proper intercalation of notochord cells.

**Figure 8:**
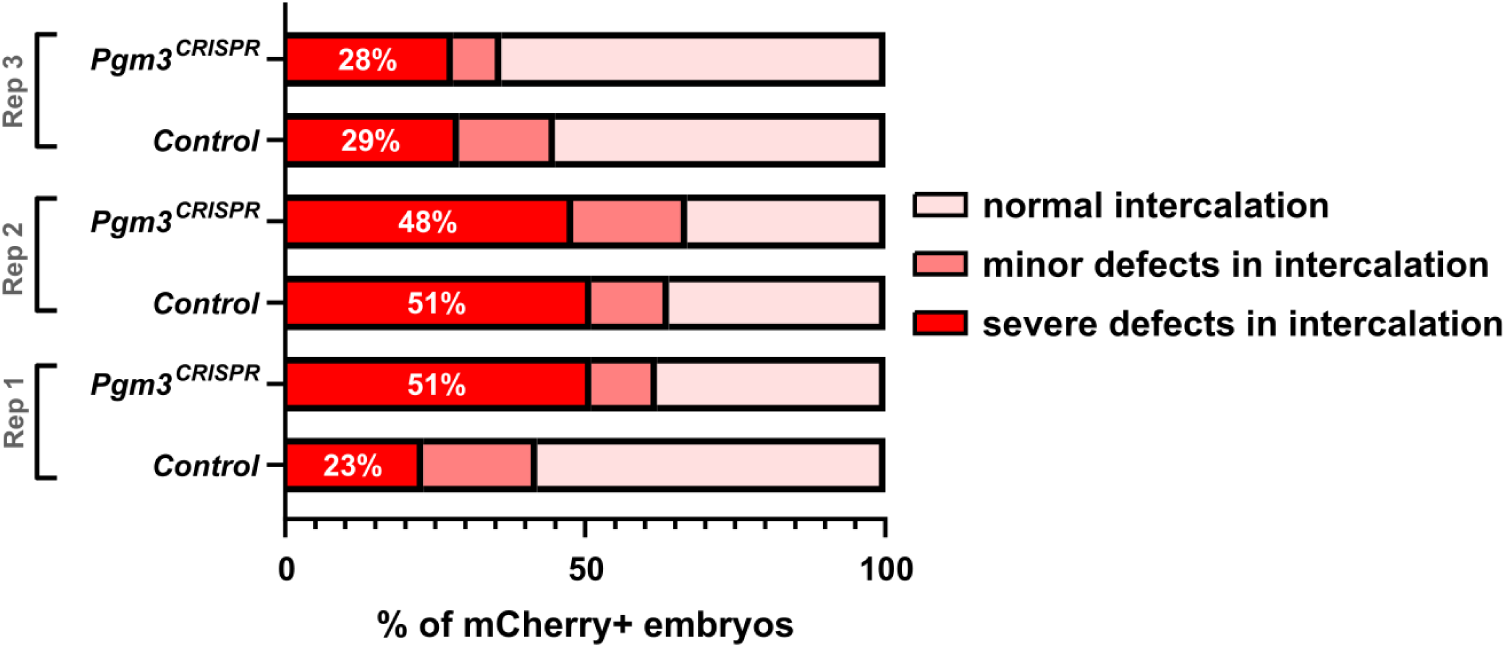
Scoring of intercalation defects in mCherry+ tailbuds in control and *Pgm3* knockout conditions in three separate replicates. Replicate 3 used Unc-76::GFP instead of CD4::GFP.. n = 75 for each condition and replicate. Control = negative control CRISPR condition.

Similarly, for *Fcho* CRISPR we performed the same intercalation assay over two replicates to determine if this gene was playing a role in this process of notochord morphogenesis. Scoring results are summarized in **Figure 9**, showing a slight increase in defects in the knockout in replicate one, though barely any increase in replicate two. Therefore we cannot conclude that Fcho is required for proper intercalation either. Finally, in a single replicate for *Nckap1* CRISPR as well, and in the first replicate, we also saw similar percentage of defects in the control vs knockout conditions, and these data are summarized in **Figure 10**.

**Figure 9:**
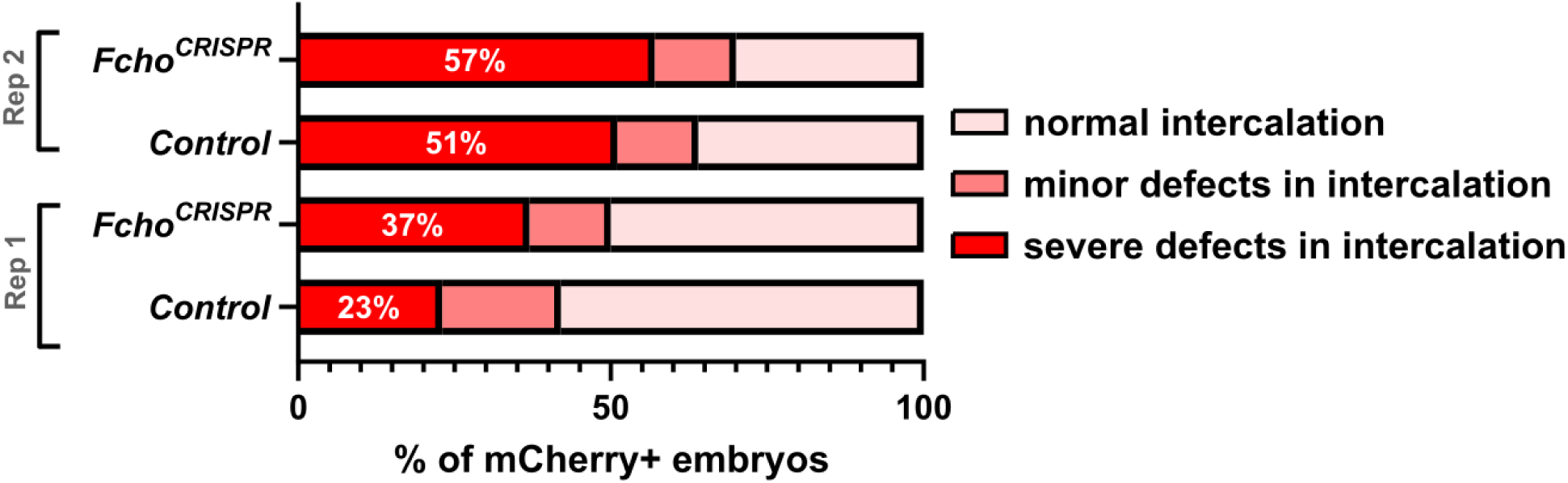
Scoring of intercalation defects in mCherry+ tailbuds in control and *Fcho* knockout conditions over two replicates. n = 75 for each condition and replicate. Control = negative control CRISPR condition.

**Figure 10:**
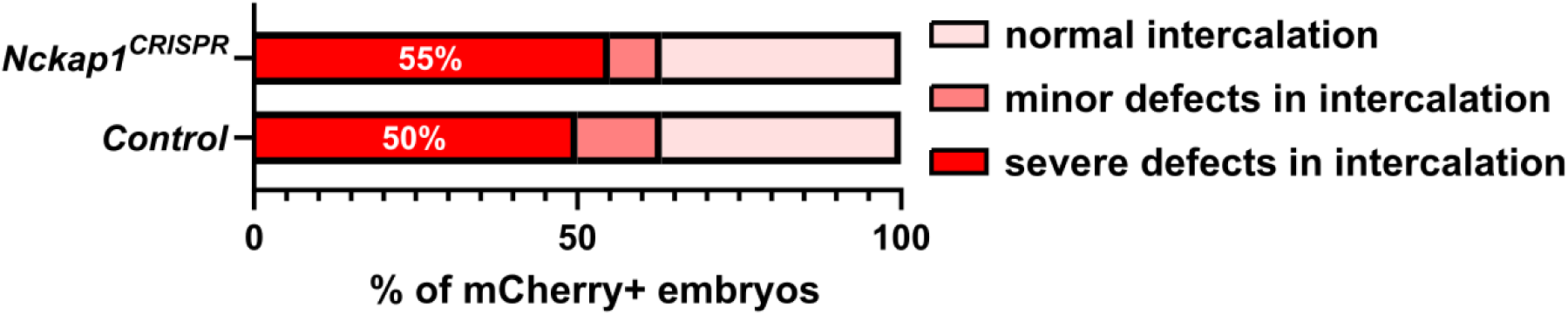
Scoring of intercalation defects in mCherry+ tailbuds in control and *Nckap1* knockout condition in one replicate. n = 75 for either condition. Control = negative control CRISPR condition.

Overall, these results do not suggest that these genes play a significant role during the early stage of intercalation in the notochord. However, we cannot rule out minor roles either. As can be seen from the variation in the incidence of severe defects among replicates, even in negative control embryos, it is clear that intercalation is a sensitive process that is especially susceptible to non-specific effects. These effects can arise from the myriad steps involved in generating tissue-specific CRISPR knockouts in F0 embryos, namely the chemical and enzymatic removal of the chorion (egg shell), or the transfection with plasmid DNAs by whole zygote electroporation. Intercalation may also be sensitive to environmental and/or seasonal conditions, as gametes are obtained from adult specimens collected weekly from various locations in the wild. Nonetheless, even with a higher baseline incidence of intercalation defects in some control samples, we would expect an increase in defects in the parallel CRISPR samples, if these genes were heavily involved in this process.

### *Pgm3* and *Fcho* are required for proper elongation of the larval tail

Next, we sought to assess the potential role of these genes in notochord elongation and tubulogenesis, using a similar tissue-specific CRISPR/Cas9-mediated mutagenesis approach as for the intercalation experiments above (Popsuj et al., 2024). Per 700 µl of electroporation volume, we used 80 µg of sgRNA plasmid (40 µg of each when combined, using same combinations as for intercalation assays), 35 µg of Cas9 plasmid, and 20 µg of *Brachyury>H2B::mCherry.* Instead of a green fluorescent reporter plasmid, we aimed to use fluorescent phalloidin to label the outline of cells to more easily assay vacuolization.

Larvae were raised to 17.5 hpf at 20°C, fixed, stained with fluorescent phalloidin (conjugated to AlexaFluor 647, from ThermoFisher), and mounted on glass slides for microscopy. Larvae were imaged using a scanning point confocal microscope (Zeiss), as well as using an epifluorescence microscope (Leica) for scoring and tail length measurements. Confocal image Z-stacks were processed in Fiji/ImageJ, while tail measurements were performed in Leica LASX software. Tails lengths were measured roughly along the anterior-posterior axis, and larvae were also qualitatively scored for wild-type or truncated tails. Raw data and images can be found at https://zenodo.org/records/18623201.

When compared to negative control larvae, *Pgm3* CRISPR and *Fcho* CRISPR larvae had visibly shortened, truncated tails. In contrast, *Nckap1* CRISPR larvae tails did not appear to be significantly different from that of negative control larvae. This was observed both through measurements of individual tail lengths (**Figure 11A**), as well as qualitative assessment of tails that appeared truncated (**Figure 11B**). Notochord cells were still arrayed more or less linearly in *Pgm3* and *Fcho* CRISPR larvae, suggesting that earlier intercalation was not affected as much as later processes of notochord cell or lumen elongation, and/or elongation of surrounding tissues such as tail muscles or endodermal strand (**Figure 12**). *Pgm3* CRISPR appeared to have the largest effect, as the reduction in tail length was most pronounced in this condition (**Figure 11A**), though qualitatively both *Pgm3* and *Fcho* CRISPR conditions were comparable as far as frequency of tail elongation defects (**Figure 11B**).

**Figure 11:**
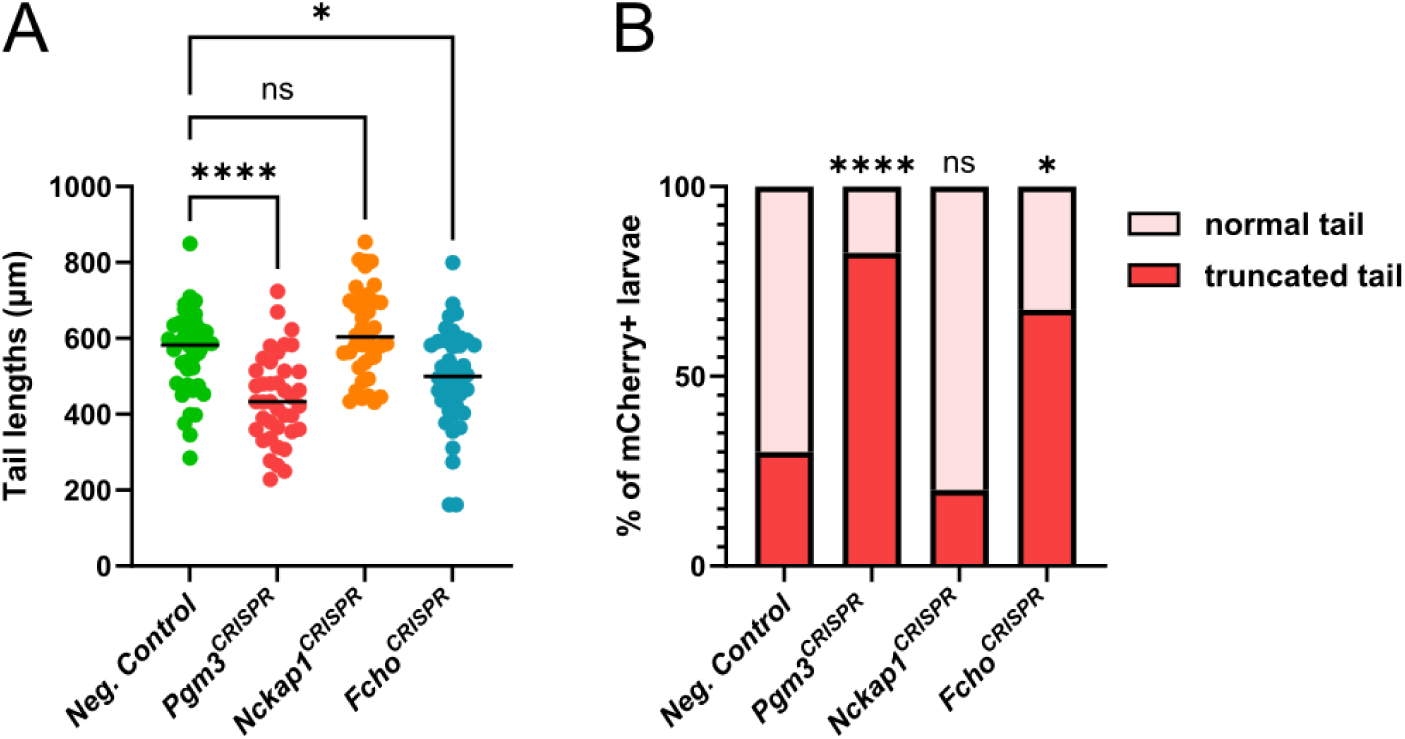
Effect of Pgm3/Nckap1/Fcho CRISPR on tail elongation. A) Plot showing tail measurements of individual larvae. n = 40 for each condition. Statistical analysis was performed as ordinary one-way ANOVA with Dunnett’s multiple comparison tests. ****: p < 0.0001, *: p = 0.0324, ns: p = 0.0789. B) Plot showing scoring same larvae as in panel A, for normal/wild type tails and for truncated tails. n = 40 for each condition. Fisher’s Exact Test was performed comparing each CRISPR condition to the negative control. ****: p < 0.0001, *: p = 0.0016, ns: p = 0.4391. Only larvae expressing Brachyury>H2B::mCherry were imaged/scored.

**Figure 12:**
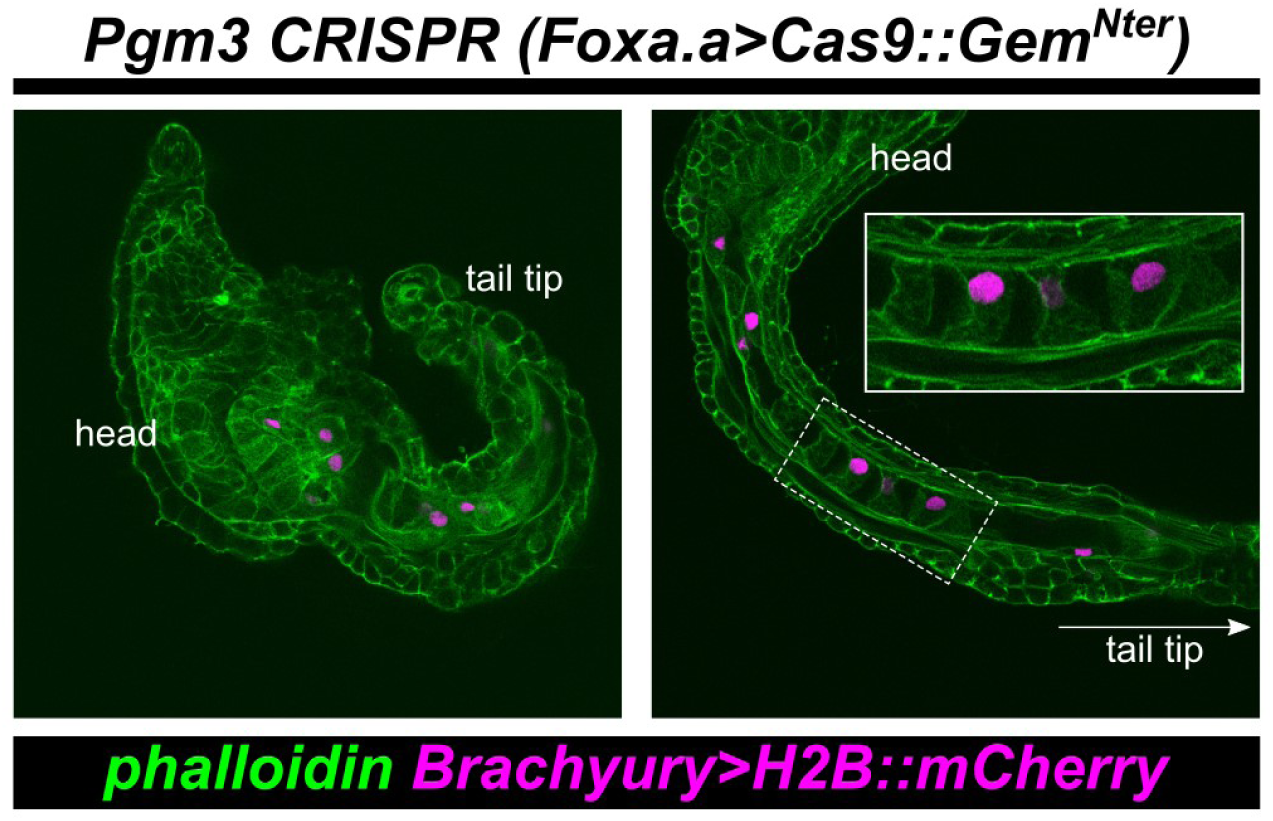
*Pgm3* CRISPR larvae. Representative confocal image slices of *Pgm3* CRISPR larvae of varying phenotype severity. Inset: notochord cells have intercalated but cell elongation and vacuole/lumen expansion is blocked. In green is AlexaFluor-647::phalloidin.

Overall, these suggest the *Pgm3* and *Fcho* might be required for tail elongation during *Ciona* embryogenesis, while *Nckap1* appears mostly dispensable (pending biological replicates). Our findings do not rule out a role for *Nckap1* in other developmental processes not assayed here, or in *Foxa.a-*negative tissues. The more pronounced effect of *Pgm3* CRISPR relative to *Fcho* CRISPR could be due to one or multiple factors, regardless of the gene products’ biological functions. *Pgm3* is the more highly and widely expressed gene according to single-cell RNA sequencing (scRNAseq) data (**Figure 1B-D**) and is upregulated in the endoderm and notochord during tail elongation according to mRNA *in situ* hybridization (Reeves et al., 2017). While the expression pattern of *Fcho* has yet to be determined by *in situs,* by either scRNAseq or bulk RNA sequencing its expression appears lower and relatively sparse (**Figure 1B-D**) Therefore, Pgm3 might be rate-limiting in more cells crucial for tail morphogenesis in *Ciona*.

Future work will be needed to determine the tissue-specific requirements for these genes, especially *Pgm3,* which is highly expressed in both endoderm and notochord. The key will be to knock out *Pgm3* only in the endoderm or the notochord and assay tail elongation. It is possible that the severe defect we describe here might be due primarily to disrupting *Pgm3* in the endoderm, or in the notochord, for instance. To do this we can use endoderm- or notochord-specific promoters to drive Cas9 expression in only one tissue at a time.

Future work is also required to elucidate the molecular mechanisms of Pgm3 and/or Fcho in tail elongation. In mammals, Pgm3 is vital for protein glycosylation. Previous work has shown that protein glycosylation is required for proper notochord lumen expansion in *Ciona* (Wang et al., 2023). We hypothesize that Pgm3 in the endoderm and/or notochord may be required for glycosylation of extracellular matrix and/or cell surface receptors involved in tail elongation and lumen expansion. Proteomic analysis of *Pgm3* CRISPR larvae might reveal such downstream glycoproteins. In fish, Fcho1 interacts with BMP receptors, and *fcho1* morphants show notochord and tail elongation defects as result of BMP signaling perturbation (Umasankar et al., 2012). BMP plays several roles in patterning the *Ciona* embryo, particularly in the tail (Christiaen et al., 2010; Lanoizelet et al., 2024; Liu et al., 2023; Pasini et al., 2006; Roure et al., 2023). Other roles for BMP or TGF-family signaling pathway in specifically notochord development and tail elongation have yet to be fully demonstrated, though Fcho could be generally affecting other receptors in an analogous manner in *Ciona*.

### Investigating a role for Plastin in papilla morphogenesis

While we were focused on knocking out *Fcho, Pgm3,* and *Nckap1* in the notochord and endoderm, we were inspired by a parallel collaboration between Arcadia and David Booth’s group at UCSF (Chou and Sun, 2025a). Zoogle identified the gene *Plastin 1 (PLS1)* as an exceptional candidate to study in the choanoflagellate *Salpingoeca rosetta* based on predicted structural similarity. This gene encodes an actin-bundling protein, and associated mutations are implicated in hearing loss in humans, likely through effects on mechanosensitive hair cell developmental defects (Morgan et al., 2019; Taylor et al., 2015).

In *Ciona,* we previously identified a *PLS1* homolog (KH gene ID KH.S521.5, KY21 gene ID KY21.Chr1.1966) as being upregulated in the developing sensory-adhesive papillae of the larva (**Figure 13**)(Johnson et al., 2024), which is a set of three protruding clusters of cells that are elongated along the apical-basal axis (Zeng et al., 2019). Of note, we found that this gene (which we call herein *Plastin*) is upregulated by the transcription factor Islet, which is crucial for the elongated cell morphology of the papillae (Johnson et al., 2024; Wagner et al., 2014). We therefore had previously hypothesized that *Ciona* Plastin might be important for regulating actin dynamics during papilla cell elongation.

**Figure 13.**
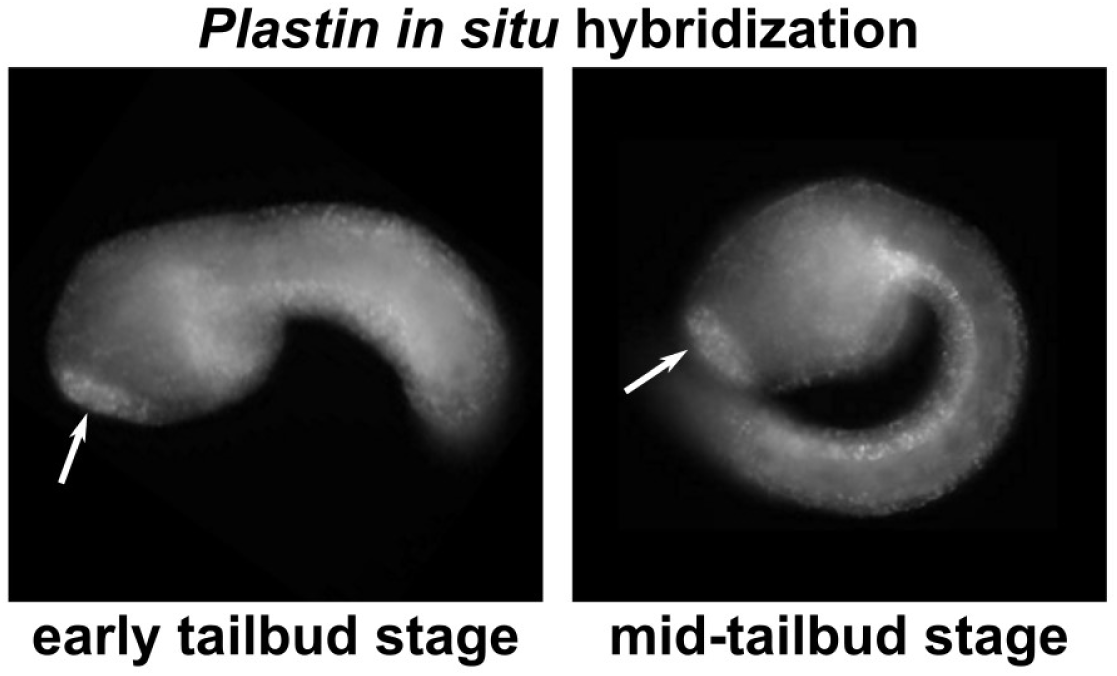
Whole-mount, fluorescent mRNA *in situ* hybridization for *Plastin* in Ciona robusta embryos, revealing strong expression in the developing papilla region (arrowheads). Expression can also be seen in the endoderm of the head/trunk and tail (endodermal strand), especially at mid-tailbud stage.

Zoogle did not suggest *Ciona* Plastin as a particularly close structural match for human Plastin 1, coming in at a very middling portfolio rank of 29 (out of 50). However, we noticed that the particular Uniprot sequence of *Ciona* Plastin used by Zoogle came from an ENSEMBL gene model that is incomplete, missing a substantial portion of the exons encoding the conserved C-terminus (**Figure 14**). We we therefore curious whether, in reality, *Ciona* Plastin might have scored higher and could serve as a good model for studying conserved Plastin functions. Here we describe our first attempt to knock out *Plastin* in the papillae of the *Ciona* larva to potentially reveal resulting defects in cell elongation.

**Figure 14.**
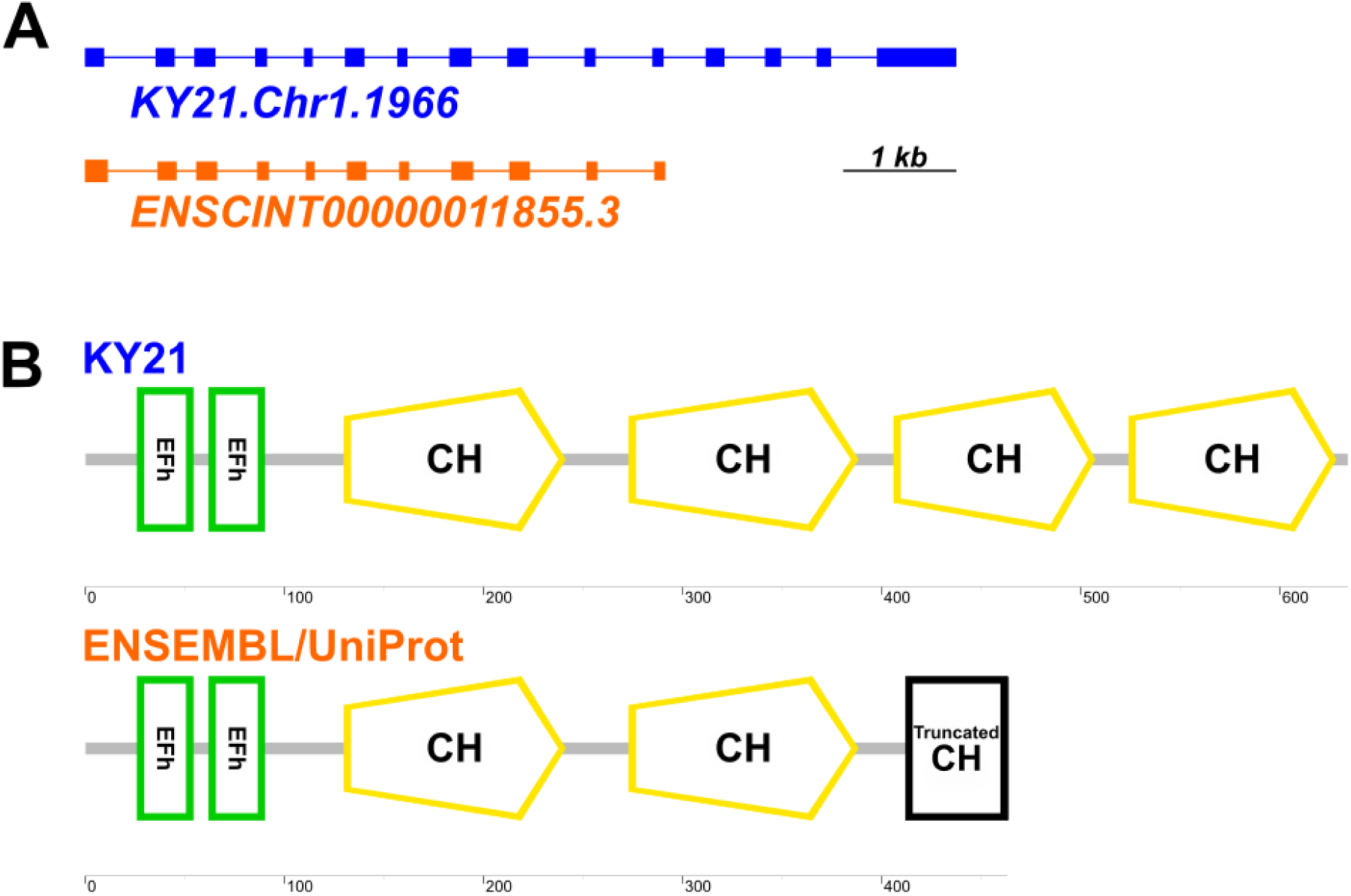
A) Diagram of KY21 (top, blue) and ENSEMBL (bottom, orange) gene models, showing that the ENSEMBL model is likely incomplete. B) Diagram comparing the predicted domain structures of the KY21 Plastin protein sequence (top) and the UniProt sequence (bottom, based on the ENSEMBL gene model). The third Calponin Homology (CH) domain is truncated, and the fourth CH domain is missing, in the UniProt sequence. EFh = EF-hand, calcium-binding motif. Protein domains analyzed by SMART (Letunic et al., 2021).

### Plastin CRISPR approach

Several single-chain guide RNAs (sgRNAs) targeting *Plastin* were designed and validated by Illumina-based amplicon sequencing followed our established routine protocol (Popsuj et al., 2024). Of these, we selected the sgRNAs shown in **Figure 15** for use in combination to knock out *Plastin* by tissue-specific CRISPR/Cas9-mediated mutagenesis. **Figure 3** shows the indel plots indicating successful targeting.

**Figure 15.**
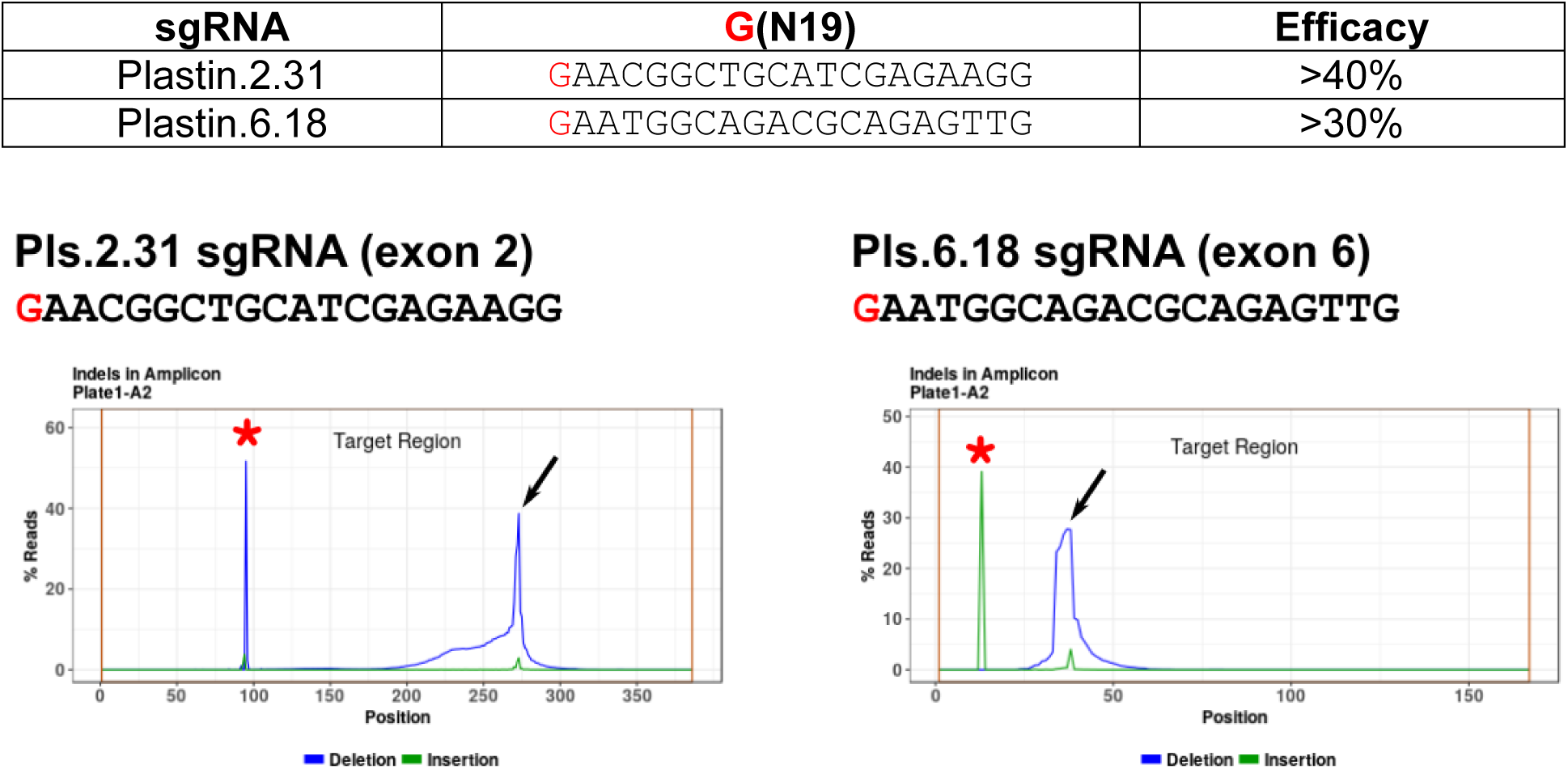
Top: *Plastin* sgRNAs selected after validation. Bottom: Insertion/deletion (indel) plots generated by Azenta/GeneWiz’s “Amplicon-EZ” illumina sequencing service. Black arrows indicate indels generated by CRISPR/Cas9 using the indicated sgRNAs. Red asterisks indicate naturally-occurring indels.

Tissue-specific CRISPR was carried out similarly as above, but using the *Foxc* promoter to drive expression of Cas9::Geminin^N-terminus^ in the papilla lineage (Johnson et al., 2024; Pennati et al., 2024). Embryos were co-electroporated with *Foxc>H2B::mCherry* reporter plasmid to label the cell nuclei of the CRISPR’d lineage, as well as a *CryBG>Unc-76::GFP* reporter plasmid to label the cytoplasm of the axial columnar cells (ACCs), the central, most elongated cell type of the papillae (Johnson et al., 2020; Shimeld et al., 2005). The electroporation recipes (per 700 µl of electroporation volume) were thus: 10 µg *Foxc>H2B::mCherry,* 35 µg *Foxc>Cas9::GemininN-terminus,* 35 µg *CryBG>Unc-76::GFP,* and 80 µg total of sgRNA plasmid (40 µg each *Plastin* sgRNA, or 80 µg of the single negative control sgRNA).

Larvae were fixed at 17 hpf (20°C) and imaged using a Leica DMI8 inverted epifluorescence microscope. Raw images can be found at https://zenodo.org/records/17892564. ACC lengths were measured along the apical-basal axis using LASX software, following the previously established method (Johnson et al., 2024). Raw lengths were plotted and analyzed in GraphPad Prism.

We observed that ACC length was not significantly reduced upon *Plastin* CRISPR in the papilla lineage (**Figure 16A,B**). This was not entirely surprising, as previous monogenic knockouts of other genes encoding cytoskeletal regulators downstream of Islet (e.g. *Villin, Tubulin alpha*) failed to result in significant, consistent shortening of the papillae (Johnson et al., 2024). This in spite of *Islet* knockouts or dominant-negatives showing significantly shortened papillae (Johnson et al., 2024; Wagner et al., 2014).

**Figure 16.**
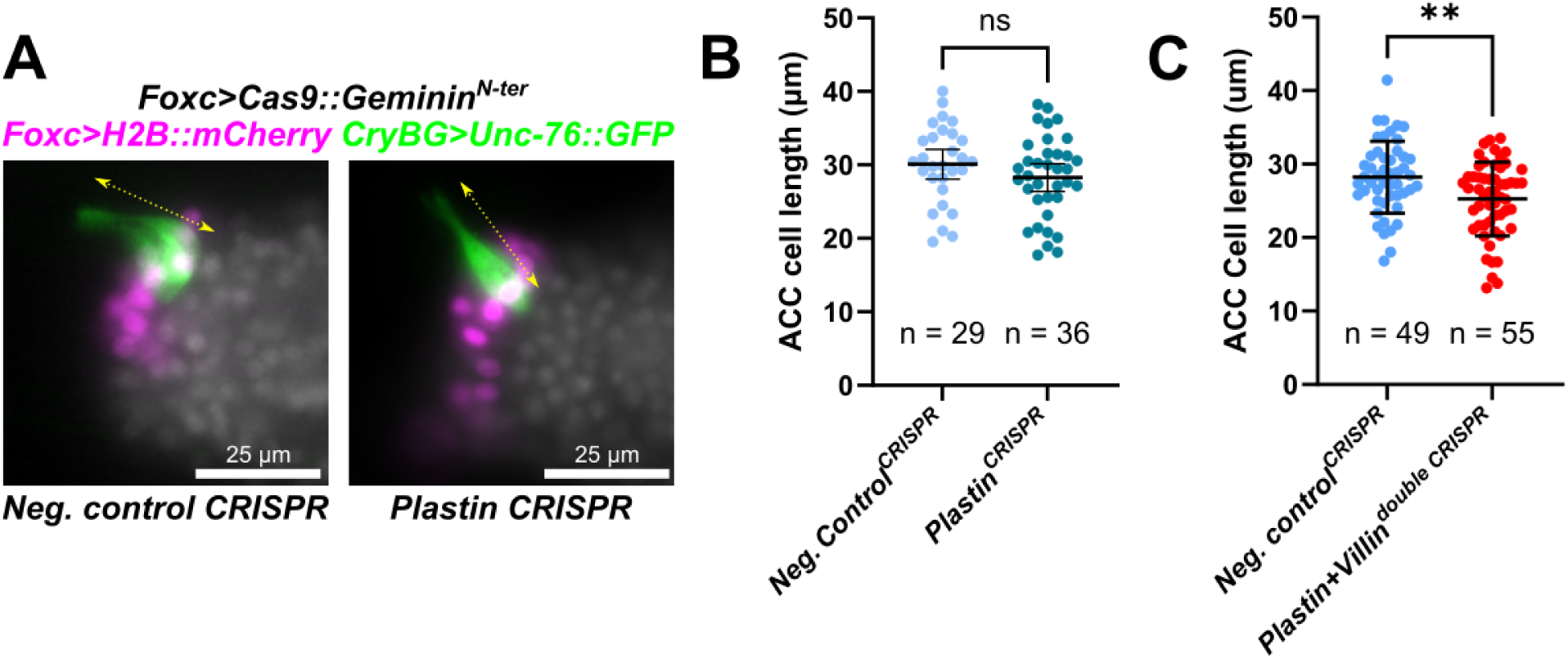
A) Images of the papillae in 17 hpf (20°C) larvae, comparing the negative control CRISPR and *Plastin* CRISPR conditions. Yellow arrow indicates axial columnar cell (ACC) length along their apical-basal axis, which was measured for this assay. Larvae counterstained with DAPI (white overlay). B) Quantification of ACC cell lengths, showing a slight but not statistically significant drop in average length upon tissue-specific *Plastin* CRISPR knockout. ns = not statistically significant, Welch’s t-test (two-tailed). C) In contrast there was a statistically significant (p = 0.0032 by two-tailed Welch’s t-test) difference in cell lengths between double *Plastin/Villin* CRISPR.

We hypothesize that multiple genes downstream of Islet may be acting in a partially overlapping or redundant manner to regulate the cytoskeleton during papilla cell elongation. For instance, *Villin* (KY21.Chr9.368) encodes another actin-bundling protein and therefore might be compensating for loss of *Plastin.* In fact, *Villin* is expressed later, specifically in elongating central *Islet+* cells that give rise to the ACCs (Johnson et al., 2024). The expression of *Plastin* during tailbud stages throughout the entire papilla territory (**Figure 14**) suggests it may be important for the earlier thickening of the papilla epithelium. Thus, it is possible that *Villin* and *Plastin* may combine in partially overlapping ways to carry out different steps in papilla morphogenesis. In mammals, a triple knockout of actin-bundling factors (Villin, Plastin1, and Epsin) did not prevent intestinal microvillus formation but reduced their length (Revenu et al., 2012).

With this in mind, we sought to perform tissue-specific, double CRISPR knockouts in the papillae of *Ciona*. For this, we used 80 g of the above *Plastin*-targeting sgRNAs combined with 35 g each of previously validated and published *Villin.4.74* and *Villin.5.105* sgRNAs (Johnson et al., 2024). When we combined both *Villin-* and *Plastin-*targeting sgRNAs (raw data accessible here: https://zenodo.org/records/21840646 ), we obtained a statistically significant reduction in ACC length compared to the negative control (**Figure 16C**). This was also in contrast to the previous attempt at knocking out *Villin* with three sgRNAs, which did not reproducibly shorten cell length (Johnson et al., 2024). Taken together, these data suggest Plastin and Villin may be acting in partially redundant ways to support cytoskeletal dynamics underlying the elongated axes of protruding papilla cells in *Ciona*.

## CONCLUSIONS

Here we have described the initial characterization of three genes in *Ciona* pinpointed by Zoogle as highly similar to their human candidate disease gene orthologs (*Fcho, Pgm3, Nckap1)*, and one that we believe may be a good match based on updated, full-length protein sequences (Plastin). Tissue-specific CRISPR knockouts of three out of these four resulted in a developmental defect. Of these three, we believe *Pgm3* to be the most promising one as its disruption had the most profound, obvious, and penetrant effect, resulting in a robust phenotype that would be amenable to higher-throughput screening. *Fcho* CRISPR resulted in a similar, but less profound defect, while an even more subtle phenotype for *Plastin* CRISPR was only unmasked by concurrent knockout of *Villin.* Using *Ciona* as a model to study Pgm3 function may be further enhanced by its potential role upstream of protein glycosylation. In the future, it may be attractive to use proteomics to identify candidate targets of Pgm3 in the *Ciona* notochord or endoderm, and devise rescue experiments aimed at identifying potential therapeutics for treating Pgm3-related human diseases. Our lab continues to use Zoogle to identify other promising disease gene candidate orthologs that may be amenable to studying in *Ciona* embryos.

## Supporting information

Supplemental Sequences File

## REFERENCES

Bernadskaya, Y., Christiaen, L., 2016. Transcriptional control of developmental cell behaviors. Annual review of cell and developmental biology 32, 77–101.

Calzoni, E., Platt, C.D., Keles, S., Kuehn, H.S., Beaussant-Cohen, S., Zhang, Y., Pazmandi, J., Lanzi, G., Pala, F., Tahiat, A., 2019. F-BAR domain only protein 1 (FCHO1) deficiency is a novel cause of combined immune deficiency in human subjects. Journal of Allergy and Clinical Immunology 143, 2317–2321.

Cao, C., Lemaire, L.A., Wang, W., Yoon, P.H., Choi, Y.A., Parsons, L.R., Matese, J.C., Levine, M., Chen, K., 2019. Comprehensive single-cell transcriptome lineages of a proto-vertebrate. Nature 571, 349–354.

Castro, C.N., Rosenzwajg, M., Carapito, R., Shahrooei, M., Konantz, M., Khan, A., Miao, Z., Groß, M., Tranchant, T., Radosavljevic, M., 2020. NCKAP1L defects lead to a novel syndrome combining immunodeficiency, lymphoproliferation, and hyperinflammation. Journal of Experimental Medicine 217, e20192275.

Chou, S., Patton, A.H., Sun, D.A., 2025. A framework for modeling human monogenic diseases by deploying organism selection. The Stacks.

Chou, S., Sun, D.A., 2025a. Modeling human monogenic diseases using the choanoflagellate Salpingoeca rosetta. The Stacks.

Chou, S., Sun, D.A., 2025b. Modeling human monogenic diseases using the tunicate Ciona intestinalis. The Stacks.

Christiaen, L., Stolfi, A., Levine, M., 2010. BMP signaling coordinates gene expression and cell migration during precardiac mesoderm development. Developmental Biology 340, 179–187.

Christiaen, L., Wagner, E., Shi, W., Levine, M., 2009a. Electroporation of Transgenic DNAs in the Sea Squirt Ciona. Cold Spring Harbor Protocols 2009, pdb.prot5345.

Christiaen, L., Wagner, E., Shi, W., Levine, M., 2009b. Isolation of Sea Squirt (Ciona) Gametes, Fertilization, Dechorionation, and Development: Figure 1. Cold Spring Harbor Protocols 2009, pdb.prot5344.

Concordet, J.-P., Haeussler, M., 2018. CRISPOR: intuitive guide selection for CRISPR/Cas9 genome editing experiments and screens. Nucleic Acids Research 46, W242–W245.

Cook, S.A., Comrie, W.A., Poli, M.C., Similuk, M., Oler, A.J., Faruqi, A.J., Kuhns, D.B., Yang, S., Vargas-Hernández, A., Carisey, A.F., 2020. HEM1 deficiency disrupts mTORC2 and F-actin control in inherited immunodysregulatory disease. Science 369, 202–207.

Corbo, J.C., Levine, M., Zeller, R.W., 1997. Characterization of a notochord-specific enhancer from the Brachyury promoter region of the ascidian, Ciona intestinalis. Development 124, 589–602.

DeWeirdt, P.C., McGee, A.V., Zheng, F., Nwolah, I., Hegde, M., Doench, J.G., 2022. Accounting for small variations in the tracrRNA sequence improves sgRNA activity predictions for CRISPR screening. Nature Communications 13, 5255.

Di Gregorio, A., 2020. The notochord gene regulatory network in chordate evolution: Conservation and divergence from Ciona to vertebrates. Elsevier, pp. 325–374.

Di Gregorio, A., Corbo, J.C., Levine, M., 2001. The regulation of forkhead/HNF-3β expression in the Ciona embryo. Developmental biology 229, 31–43.

Doench, J.G., Fusi, N., Sullender, M., Hegde, M., Vaimberg, E.W., Donovan, K.F., Smith, I., Tothova, Z., Wilen, C., Orchard, R., 2016. Optimized sgRNA design to maximize activity and minimize off-target effects of CRISPR-Cas9. Nature biotechnology 34, 184–191.

Greig, K.T., Antonchuk, J., Metcalf, D., Morgan, P.O., Krebs, D.L., Zhang, J.-G., Hacking, D.F., Bode, L., Robb, L., Kranz, C., 2007. Agm1/Pgm3-mediated sugar nucleotide synthesis is essential for hematopoiesis and development. Molecular and cellular biology 27, 5849–5859.

Henne, W.M., Boucrot, E., Meinecke, M., Evergren, E., Vallis, Y., Mittal, R., McMahon, H.T., 2010. FCHo proteins are nucleators of clathrin-mediated endocytosis. Science 328, 1281–1284.

Hernandez, S.A., Johnson, C.J., Stolfi, A., 2025. Using CRISPR/Cas9 to investigate the developmental roles of Fcho, Pgm3, and Nckap1 in Ciona, Part 2: testing their roles in notochord intercalation and convergent extension. Zenodo, https://zenodo.org/records/17880460.

Hernandez, S.A., Stolfi, A., 2026. Using CRISPR/Cas9 to investigate the developmental roles of Fcho, Pgm3, and Nckap1 in Ciona, Part 3: testing their roles in tail morphogenesis. Zenodo, https://zenodo.org/records/18623412.

Hsu, P.D., Scott, D.A., Weinstein, J.A., Ran, F.A., Konermann, S., Agarwala, V., Li, Y., Fine, E.J., Wu, X., Shalem, O., 2013. DNA targeting specificity of RNA-guided Cas9 nucleases. Nature biotechnology 31, 827–832.

Jiang, D., Smith, W.C., 2007. Ascidian notochord morphogenesis. Developmental Dynamics 236, 1748–1757.

Johnson, C.J., Hernandez, S., Alberto, S., Using CRISPR/Cas9 to investigate the developmental roles of Fcho, Pgm3, and Nckap1 in Ciona, Part 1: guide RNA design and validation.

Johnson, C.J., Hernandez, S.A., Stolfi, A., 2025. Using CRISPR/Cas9 to investigate the developmental roles of Fcho, Pgm3, and Nckap1 in Ciona, Part 1: guide RNA design and validation. Zenodo, https://zenodo.org/records/17095188.

Johnson, C.J., Kulkarni, A., Buxton, W.J., Hui, T.Y., Kayastha, A., Khoja, A.A., Leandre, J., Mehta, V.V., Ostrowski, L., Pareizs, E.G., Scotto, R.L., Vargas, V., Vellingiri, R.M., Verzino, G., Vohra, R., Wakade, S.C., Winkeljohn, V.M., Winkeljohn, V.M., Rotterman, T.M., Stolfi, A., 2023. Using CRISPR/Cas9 to identify genes required for mechanosensory neuron development and function. Biology Open 12.

Johnson, C.J., Razy-Krajka, F., Stolfi, A., 2020. Expression of smooth muscle-like effectors and core cardiomyocyte regulators in the contractile papillae of Ciona. EvoDevo 11, 1–18.

Johnson, C.J., Razy-Krajka, F., Zeng, F., Piekarz, K.M., Biliya, S., Rothbächer, U., Stolfi, A., 2024. Specification of distinct cell types in a sensory-adhesive organ important for metamorphosis in tunicate larvae. PLoS biology 22, e3002555.

Johnson, C.J., Stolfi, A., 2025. Using CRISPR/Cas9 to investigate the developmental roles of Plastin in Ciona, Part 1: testing its role in papilla cell elongation. Zenodo, https://zenodo.org/records/17904585.

Lanoizelet, M., Elkhoury Youhanna, C., Roure, A., Darras, S., 2024. Molecular control of cellulosic fin morphogenesis in ascidians. Bmc Biology 22, 74.

Letunic, I., Khedkar, S., Bork, P., 2021. SMART: recent updates, new developments and status in 2020. Nucleic acids research 49, D458–D460.

Liu, B., Ren, X., Satou, Y., 2023. BMP signaling is required to form the anterior neural plate border in ascidian embryos. Development Genes and Evolution, 1-11.

Łyszkiewicz, M., Ziętara, N., Frey, L., Pannicke, U., Stern, M., Liu, Y., Fan, Y., Puchałka, J., Hollizeck, S., Somekh, I., 2020. Human FCHO1 deficiency reveals role for clathrin-mediated endocytosis in development and function of T cells. Nature communications 11, 1031.

Morgan, A., Koboldt, D.C., Barrie, E.S., Crist, E.R., García García, G., Mezzavilla, M., Faletra, F., Mihalic Mosher, T., Wilson, R.K., Blanchet, C., 2019. Mutations in PLS1, encoding fimbrin, cause autosomal dominant nonsyndromic hearing loss. Human mutation 40, 2286–2295.

Nydam, M.L., Harrison, R.G., 2010. Polymorphism and divergence within the ascidian genus Ciona. Molecular Phylogenetics and Evolution 56, 718–726.

Park, H., Staehling-Hampton, K., Appleby, M.W., Brunkow, M.E., Habib, T., Zhang, Y., Ramsdell, F., Liggitt, H.D., Freie, B., Tsang, M., 2008. A point mutation in the murine Hem1 gene reveals an essential role for Hematopoietic protein 1 in lymphopoiesis and innate immunity. The Journal of experimental medicine 205, 2899–2913.

Pasini, A., Amiel, A., Rothbächer, U., Roure, A., Lemaire, P., Darras, S., 2006. Formation of the ascidian epidermal sensory neurons: insights into the origin of the chordate peripheral nervous system. PLoS biology 4, e225.

Pennati, A., Jakobi, M., Zeng, F., Ciampa, L., Rothbächer, U., 2024. Optimizing CRISPR/Cas9 approaches in the polymorphic tunicate Ciona intestinalis. Developmental Biology 510, 31–39.

Popsuj, S., Cohen, L., Ward, S., Lewis, A., Yoshida, S., Herrera, R.A., Cota, C.D., Stolfi, A., 2024. CRISPR/Cas9 protocols for disrupting gene function in the non-vertebrate chordate Ciona. Integrative and Comparative Biology 64, 1182–1193.

Popsuj, S., Di Gregorio, A., Swalla, B.J., Stolfi, A., 2023. Loss of collagen gene expression in the notochord of the tailless tunicate Molgula occulta. Integrative and Comparative Biology, icad071.

Popsuj, S., Kalsang, T., Kim, K., Drummond, E., Manekar, P., Munagapati, P., Oleti, M., Sato, H., Vickery, I., Gigante, E.D., 2026. Validated CRISPR/Cas9 guide RNAs targeting neurodevelopmental genes in the tunicate Ciona robusta. Differentiation, 100973.

Reeves, W.M., Wu, Y., Harder, M.J., Veeman, M.T., 2017. Functional and evolutionary insights from the Ciona notochord transcriptome. Development 144, 3375–3387.

Revenu, C., Ubelmann, F., Hurbain, I., El-Marjou, F., Dingli, F., Loew, D., Delacour, D., Gilet, J., Brot-Laroche, E., Rivero, F., Louvard, D., Robine, S., 2012. A new role for the architecture of microvillar actin bundles in apical retention of membrane proteins. Mol Biol Cell 23, 324–336.

Roure, A., Chowdhury, R., Darras, S., 2023. Regulation of anterior neurectoderm specification and differentiation by BMP signaling in ascidians. Development 150.

Satoh, N., 2013. Developmental genomics of ascidians. John Wiley & Sons.

Satou, Y., Nakamura, R., Yu, D., Yoshida, R., Hamada, M., Fujie, M., Hisata, K., Takeda, H., Satoh, N., 2019. A nearly complete genome of Ciona intestinalis Type A (C. robusta) reveals the contribution of inversion to chromosomal evolution in the genus Ciona. Genome biology and evolution 11, 3144–3157.

Shimeld, S.M., Purkiss, A.G., Dirks, R.P.H., Bateman, O.A., Slingsby, C., Lubsen, N.H., 2005. Urochordate βγ-crystallin and the evolutionary origin of the vertebrate eye lens. Current biology 15, 1684–1689.

Stray-Pedersen, A., Backe, P.H., Sorte, H.S., Mørkrid, L., Chokshi, N.Y., Erichsen, H.C., Gambin, T., Elgstøen, K.B.P., Bjørås, M., Wlodarski, M.W., 2014. PGM3 mutations cause a congenital disorder of glycosylation with severe immunodeficiency and skeletal dysplasia. The American Journal of Human Genetics 95, 96–107.

Taylor, R., Bullen, A., Johnson, S.L., Grimm-Günter, E.-M., Rivero, F., Marcotti, W., Forge, A., Daudet, N., 2015. Absence of plastin 1 causes abnormal maintenance of hair cell stereocilia and a moderate form of hearing loss in mice. Human Molecular Genetics 24, 37–49.

Umasankar, P.K., Sanker, S., Thieman, J.R., Chakraborty, S., Wendland, B., Tsang, M., Traub, L.M., 2012. Distinct and separable activities of the endocytic clathrin-coat components Fcho1/2 and AP-2 in developmental patterning. Nature cell biology 14, 488–501.

Wagner, E., Stolfi, A., Choi, Y.G., Levine, M., 2014. Islet is a key determinant of ascidian palp morphogenesis. Development 141, 3084–3092.

Wang, Z., Tan, Z., Bi, J., Liu, A., Jiang, A., Dong, B., 2023. Proteomic identification of intracellular vesicle trafficking and protein glycosylation requirements for lumen inflation in Ciona notochord. Proteomics 23, 2200460.

Winslow, A., Jalazo, E.R., Evans, A., Winstead, M., Moran, T., 2022. A De Novo cause of PGM3 deficiency treated with hematopoietic stem cell transplantation. Journal of Clinical Immunology 42, 691–694.

Yang, L., Zerbato, B., Pessina, A., Brambilla, L., Andreani, V., Frey-Jakobs, S., Fliegauf, M., Barbouche, M.-R., Zhang, Q., Chiaradonna, F., 2024. PGM3 insufficiency: a glycosylation disorder causing a notable T cell defect. Frontiers in immunology 15, 1500381.

Zeng, F., Wunderer, J., Salvenmoser, W., Hess, M.W., Ladurner, P., Rothbächer, U., 2019. Papillae revisited and the nature of the adhesive secreting collocytes. Developmental biology 448(2), 183–198.

