## Supplemental Sequences File for "Using CRISPR/Cas9 to investigate the role of candidate human disease gene orthologs in Ciona"

**Supplemental sequences – Hernandez, Johnson et al. 2026**

**Fcho, Pgm3, and Nckap1 sgRNAs tested (g+N19)**

**Fcho.1.34	gGCACCGTATCCTGGACTGT**

**Fcho.4.63	gAAATTATCCAGCTGTCACA**

**Fcho.5.112	gGCTTTGTATGGAGACTGAA**

**Pgm3.1.83	gGCATTACCTCTAAAACCAG**

**Pgm3.2.27	gGGTAAGCATTTCACCATGA**

**Pgm3.3.31	gTAAGGCCGTGGCTCCATCA**

**Nckap1.2.74	gTGTCAGCATGCCAACAGCA**

**Nckap1.7.36	gCGTCTTGGTCAAATGATCG**

**Nckap1.8.103	gCAGGGTTAAGCATCTGTGC**

Validation primer (forward) Fcho.1.34:	GACCAACATTTAGGCCATAATG

Validation primer (reverse) Fcho.1.34:	TGGTACTGCAGTGATGTAGTG

Validation primer (forward) Fcho.4.63:	AGTTTGGACCATGAGTCAATG

Validation primer (reverse) Fcho.4.63:	AATCACCACAATTTAGAGAGTTG

Validation primer (forward) Fcho.5.112:	ACAAGTTCAAATGACACAAAGTAC

Validation primer (reverse) Fcho.5.112:	TTCCGATATCATTTCCGTACTAG

Validation primer (forward) Pgm3.1.83:	TTTGAACTGCGGCGTTTTAC

Validation primer (reverse) Pgm3.1.83:	CTCTGGGTTATGGGAAGCAG

Validation primer (forward) Pgm3.2.27:	ACACAGGATGTATGACTGTTG

Validation primer (reverse) Pgm3.2.27:	TCCTACCTCGTGTCTCTTG

Validation primer (forward) Pgm3.3.31:	TCGGTTGTGATTTAGAGCAC

Validation primer (reverse) Pgm3.3.31:	AAAGGACCAACCTCATTTAGT

Validation primer (forward) Nckap1.2.74:	ACATTGCTTTGCTCCGTAATC

Validation primer (reverse) Nckap1.2.74:	GCTTTGATCTGCTGTCACTG

Validation primer (forward) Nckap1.7.36:	ACCAAGTATGTATAGTGCATCAC

Validation primer (reverse) Nckap1.7.36:	TGAAATGTTAAATCACATTACCCAT

Validation primer (forward) Nckap1.8.103:	CTTTACTTGTATTTGCAGTCCATTC

Validation primer (reverse) Nckap1.8.103:	ATAAAGTCACGACCCGTATAC

**Plastin sgRNAs validated and selected (g+N19)**

**Plastin.2.31	gAACGGCTGCATCGAGAAGG**

**Plastin.6.18	gAATGGCAGACGCAGAGTTG**

Validation primer (forward) Plastin.2.31:	ATAAAGTGTACAAGATAGGACATC

Validation primer (reverse) Plastin.2.31:	GTATTACCCGGGCAAATTC

Validation primer (forward) Plastin.6.18:	ATGTAAGCAAATCGTGAAAAGGT

Validation primer (reverse) Plastin.6.18:	ATTTATACCTGGGTTGTGAATC

**Published Villin sgRNAs from Johnson et al. 2024 (g+N19)**

**Villin.4.74	gTTCAATACAGAGAAACACA**

**Villin.5.105	gGTCCAAGCAAAGTCCTGCT**

>Fcho protein sequence (translated from KY21.Chr8.771.v1.nonSL2-2)

MVDLLDGFTFGPTVQDTVPDDDYFAQKFWGEKNNGFDVLYHNMKHGQISVKSLQDLFRESATTEDVYAKHLTKLAKQAANSSPLGSFAPMFEVVKVMSEKLSSCHMDCVHKLHEIIKELAKYLEEQKNKHKQVKDELSGTADALHIIQTTTAAVNKSKEKYHQLCMETERLRRASAPNKEIEKAENKVKKSAEEYRAIVAKYATVRSDFEQKMTDSAKRFQEIEEAHLRHMKGVLDNYILSFENANVLINQVHQEFRRQVDEKTIASLIKQYSDTKGTGTEKPGLMVFEEWDPTTLVSIETSSTNGTLHEETQNPAPAKTRRPMALPFRRKKQRKSQKDMMNGSNTSKDTGTDKDGESIESPDKDPSVKIDEEGYTIRPETPKGSKNSWGMESSGSSDSDSDDYVSRKIHVKINPKDVTRTESGNNLDTLNAITRNLTLNSPTPVGGKRLSPLNYNSKKNKDSPAADKQEKSMSDELLELFGGGSMSSAPPPPRKNSSKADSILELYSTKPDSPATAAKKDHLTPTPIPQSSSNVSLNNLFIMPEKDAEIVTSSPATSNHNSDHLSPSHNSVDSIELNTPEQTERPRNLSLTTAWDDLGKGTPMDSFASSRGPSPLTLMHGDPIPIAVAFTETVNAYFKGSDETRCMVKITGEVQMSFPAGIVRAFTSNPNPATLSFKVKGVSNIKDFAPNANLIFADGSELSAESRSFFFNMSALVNHVKKMTEANPNSPYYNIQVMSYQVTSGYDMVPLHVTSSWKCEKDSTDVIVDYKYNTKLHLPPLKNVQIIVPVDGGVTSVQSVPTANWMSAQNRLSWKMPVPVSSMSAPTHLKSKLSLSNGPSRASALAVQFMSDGATLTGADFELTGAGYRLSLVKRKFVSGKYLSDGS*

>Pgm3 protein sequence (translated from KY21.Chr1.427.v1.SL2-1)

MATLNEIPILKLAEVHAKPKGHQMSYGTAGFRGNASGMDHVFFRMGMLAVLRSKLTEATIGVMVTASHNPEHDNGVKLIDPHGEMLTQSWEELATSLANVTNDELIKEMKKIVDSQNIQLSNEASVFIARDTRPSSLALSQAVLDGATALGATCTNYGLLTTPQLHHIVACYNGGNHNVNEEQYYQHYANAFKALLNEKPVGSSVTVDCANGVGAPKLVKLAEHIGVNIIDIVVHNNGQSGKLNENCGADYVKVQQRAPVGLNMEPDHRYASFDGDADRLVYYTLDSDCNFVLLDGDKIAALFAVYIKELLNKADINVRLGVVQTAYANGSSTNYISTEENIEVACAQTGVKHLHKVATAFDIGVYFEANGHGTVTVKDDCLNKIRSAATSEEKLQEQKLAANKLTAFLDVVNQTVGDAMSDLLAVEAILQDKGWSIRDWSCIYTELPNRQAKVKVADRTVIETTDAERKVTKPPALQPAIDKVVAEYPCGRSFVRPSGTEDVVRVYAESDTQENTDKLAKRVSLLVYELAGGVGEKPQ*

>Nckap1 protein sequence

MSISVTSRSIMARVVPGQQKLAEKLIILNNRAVGMLTRIYNIKKACSDSRSKPTYLTDKALESGIKFIVRKFPQVDTRNNQNQLAGVNNAKNEILKGLSLYYHTFVDVLEFKDHFYELLATFDSLSIYLDLTVNYDLTSHFLELIVRYASLMILVSRIDDRKAIIGLYDHAHDMIHGKMEKDYARLGQMIVDFENPLKKLVEDFGAHSRSIQNAVMSLLQIYPQRNSGADVWRSSSLFSVIATPAQMLNPAVTDKMQCEYISVESLERWIYFGYLLCHTSMVSSQEATAGLWRPVLQSNFCLTLFRDEVIMLHRSAEEVFSSIKGYNKKISEVKECRDMAAQQAGKIHKDKRKYLRSSMKEFVAVLTDQPGLLGPKTLFVFMALSYARDEVMWLARNSSNFIKKVNAEDFVDRHLGELLFYMEEIRHLVRRYDEVVQRYYVQYMYGYDSVILNEAVQNLSVCPEDESVIMTSIVNTMTSLSIKQVEQKELFDFRAMRLDWFRLQAYTSVNKAALSLKDNVVLARLMNTISFHCKVVDEIDEVLHETSDLSNFCFYQNFYATAFKRCIELPAQSRYSIIFPLICGHFVTAVGDFCPEERSHVRDRALNCVNQFLEEIAKEARNLLFNIASEQSQLAENLLPKNAAVMMKQGLVKKGKGKKHTKGNVQETTKPGLESKRKDRLYVTRMDKYHMALAEVCSAINYRANFVVWEHTFAPKEYLTAHLESRFAKHLVSMTFNKETSEIAKPSELLCKLRAYMATLQTVENYVHLDVTRIFNSVLLQQSQMVDSHGETTITTLYTQWYLEVMLRQVSGGSTVFSEMRRHFVKVPNYADSSSSNQPIINAEEYSNINELKALAQIIGPYGMKFLNESLVWHIASQITELKKSVVDNMDTLTGLRTNFDKPEQMAMLYRKLEGVENFLLRMTIIGVIFSFRDVAQEALNDVLQHRIPFLLASIADFKEHIPKETDIKVTININELASAAGIPCEIDPTLCAALSAQKIENPDEEYKVACLLMVFLAVDIPVLARNERSVFLPELVAHGNNCHCLARAVNHVAAALFTVHRGTVEDRLKEFLALASSSLLKLGQETDKVSTRSRESVYLLLERIIHFSPFLTMDLLESCFPYVLLRNSYHAVRRNQPAMTGLSRNESISATVA*

>Plastin *in situ* hybridization probe template sequenceATGGCAAATGATGAATATGAATCCGGAATCAGACAAGCATTGGCAAATATGTCACTTGGAACAGATGATGTGGATACTGTTGTTAATGCATTTATAACTGTTGACACCAACCAGAACGGCTGCATCGAGAAGGAGGAAGTAGAAGACTTGTTTAGAGAGTCGGGTTTAAAACTTCCAAAGTATAAGATCCGTGATTTAATGAAGGAAGCTGATTTGGATCGAAACAATACCATCAGCCCAACAGAATTTGCCCGGATCTACGCCCAGCTGACATCTGAAGAATATTCCCGTCAATTTAAAACTGCAATCACTACAAAGTCAAACTTGAAAAAATCAGAAATATCCAATGCGTCTGCTGAAGGGACTCAACATTCTTATTCGGAAGAAGAATGTGCTGCGTTCAACAAATGGATCACAAAGAATTTGAAAGACGATGATGATTGTAAAGATCGAGTGAAGAACATGCAAGCTAACGATTTCTTTAAACGGATGAAAGATGGAATTATTCTATGTAAGATGGTAAATCTTTCCCAACCAGACACCATTGATGAGAGAACAATAAATAAGAAAAACCTTAACATTTATCGAGAACAGGAAAATATCAATCTTGCCCTCAACTCTGCGTCTGCCATTGGATGCAACATTGTGAACATAGGAGCCGAGGATATTGTTCAAAAGAAAGAGCATCTCATCCTCGGTCTTTTATGGCAAGTCATCAGAATCGGACTTTTCAGAAAGATCGATTTGATTCACAACCCAGGTATTAGTGCTCTTCTGTTAGAAGGTGAAACATTGGATGACCTTAGGGCAATGTCACCTGAAGATCTCCTTTTAAGGTGGATGAATTATCATCTTCAACAATCTGACAAATACAAAGAGGCAACTGGAGGAAAAGTTATCACAAACTTCAGCGCAGACATTAAGGACTCCATTGCTTACACTTGTCTATTGGAGAGAATCCAACCAATTGATGAAGAGACACAACAATATGAACTGTCTCCACCAATTTGTGCCAAAATTGAAGCAAAATCCAACACTGCCAGAGCTGAATCCATGCTTGAAGATGCCGAGCGAATGAATTGCCGAGAGTTTGTGACCGCTAAAGACATTACCAAAGGAAATGCTAAACTCAACATGGCTTTCGTGGCAAACCTATTCAACACACACCCTGCCCTTACACCAAGAGACATGGAAGAGATAGAAGAGGAAAAGAGAGAAGTGAAGACATATAGAAACTGGATGAACAGTTTGGGAGTTTCTCCTCGTGTCAACAAGTTCACCAGAGACTTAACTTCTGGACTTGTTCTGTTTCAACTATACGAGCAAGTTAAGCCAGGTTGCGTGGATTGGTCCAAGGTTGATAAAAAGTGCAAAGGAAGATTCAAAAAACT

>KY21.Chr1.1966.v1.SL1-1 gene model protein sequence for Plastin

missing in ENSEMBL/UniProt model

MANDEYESGIRQALANMSLGTDDVDTVVNAFITVDTNQNGCIEKEEVEDLFRESGLKLPKYKIRDLMKEADLDRNNTISPTEFARIYAQLTSEEYSRQFKTAITTKSNLKKSEISNASAEGTQHSYSEEECAAFNKWITKNLKDDDDCKDRVKNMQANDFFKRMKDGIILCKMVNLSQPDTIDERTINKKNLNIYREQENINLALNSASAIGCNIVNIGAEDIVQKKEHLILGLLWQVIRIGLFRKIDLIHNPGISALLLEGETLDDLRAMSPEDLLLRWMNYHLQQSDKYKEATGGKVITNFSADIKDSIAYTCLLERIQPIDEETQQYELSPPICAKIEAKSNTARAESMLEDAERMNCREFVTAKDITKGNAKLNMAFVANLFNTHPALTPRDMEEIEEEKREVKTYRNWMNSLGVSPRVNKFTRDLTSGLVLFQLYEQVKPGCVDWSKVDKKCKGRFQKLQNLQYAVEVAKQLGFVIVGIEGNDILAENETLVLAVVWQIMRAYTFKILETLSEDGKPVKEQIIVDWVNQKLSDSGKETQISSFKDSEIKKSLVVIDLIDAIVPGSIRYEVVTAGETEEDQYSNAKYAVSMARKIGARIYALPDDLVEGKAKMVATIFACLMGRGMKEVE

>ENSEMBL/UniProt gene model protein sequence for Plastin

MANDEYESGIRQALANMSLGTDDVDTVVNAFITVDTNQNGCIEKEEVEDLFRESGLKLPKYKIRDLMKEADLDRNNTISPTEFARIYAQLTSEEYSRQFKTAITTKSNLKKSEISNASAEGTQHSYSEEECAAFNKWITKNLKDDDDCKDRVKNMQANDFFKRMKDGIILCKMVNLSQPDTIDERTINKKNLNIYREQENINLALNSASAIGCNIVNIGAEDIVQKKEHLILGLLWQVIRIGLFRKIDLILEPGISALLLEGETLDDLRAMSPEDLLLRWMNYHLQQSDKYKEATGGKVITNFSADIKDSIAYTCLLERIQPIDEETQQYELSPPICAKIEAKSNTARAESMLEDAERMNCREFVTAKDITKGNAKLNMAFVANLFNTHPALTPRDMEEIEDSKLSPPTEEKREVKTYRNWMNSLGVSPRVNKFTRDLTSGLVLFQLYEQVK
